# Homocitrullinated Peptides Drive Pro-Inflammatory T-Cell Responses in a Humanized Mouse Model of Rheumatoid Arthritis

**DOI:** 10.1101/2025.10.30.685402

**Authors:** Sofya Ulanova, Jaspreet Kaur, S.M. Mansour Haeryfar, Lisa Cameron, Ewa Cairns, Lillian Barra

**Affiliations:** Department of Microbiology & Immunology, Western University, London, Ontario, Canada; Department of Medicine, Division of Clinical Immunology and Allergy, Western University, London, Ontario, Canada; Department of Surgery, Division of General Surgery, Western University, London, Ontario, Canada; Department of Oncology, Western University, London, Ontario, Canada; Department of Pathology and Laboratory Medicine, Western University, London, Ontario, Canada; Department of Medicine, Division of Rheumatology, Western University, London, Ontario, Canada; Department of Epidemiology and Biostatistics, Western University, London, Ontario, Canada

**Keywords:** Rheumatoid arthritis, homocitrullinated peptides, CD4+ T cells, CD8+ T cells, shared epitope, HLA-DR4, humanized mice, autoimmunity, inflammation, cytokines

## Abstract

**Objective:** Anti-homocitrullinated protein/peptide antibodies (AHCPA) are specific to rheumatoid arthritis (RA) and predictive of worse prognosis, suggesting a pathogenic role for autoreactivity to homocitrullinated antigens. However, T-cell responses to homocitrullinated peptides remain largely unexplored. We investigated these responses in a humanized HLA-DR4-transgenic (DR4tg) mouse model of RA, which expresses the strongest genetic risk factor for this disease.

**Methods:** DR4tg mice were injected subcutaneously with a homocitrullinated peptide called HomoCitJED while control mice received phosphate-buffered saline. After 10 days, T-cells were analyzed for their phenotypic characteristics, cytokine production, and proliferative capacities in the draining lymph nodes (dLNs) and spleens by flow cytometry, enzyme-linked immunosorbent assays, and ProQuantum™ immunoassays.

**Results:** HomoCitJED immunization drove robust expansion of T helper (Th) 1, Th17 and hybrid Th1/Th17 CD4+ T cells in dLNs, alongside elevated CD25 activation marker within these subsets. Intracellular cytokine staining confirmed effector activity, revealing higher frequencies of IL-17A+, TNF-α+IL-17A+, and IFN-γ+IL-17A+ CD4+ T cells. CD4+ T cells from dLNs also up-regulated the exhaustion markers LAG-3 and Tim-3. CD8+ T cells (Tc) mirrored these findings, as HomoCitJED immunization augmented CD25 and KLRG1 expression, Tc1/Tc17-type IL-17A and IFN-γ/IL-17A production, and antigen-specific proliferation. Additionally, Tim-3, LAG-3 and PD-1 expression on these subsets was augmented.

**Conclusions:** A homocitrullinated peptide elicited Th1/Th17 and Tc1/Tc17 responses marked by concurrent activation and exhaustion signatures, pointing to a dysregulated T-cell state. These findings position homocitrulline-driven T-cell imbalance across both CD4+ and CD8+ T-cell compartments as a potential mechanistic contributor to RA pathogenesis and a target for future immunomodulatory therapies.

## Introduction

Rheumatoid arthritis (RA) is a chronic systemic autoimmune disease characterized by persistent synovial inflammation, leading to joint damage [1]. Although the etiology of RA remains unclear, one hallmark is the production of autoantibodies targeting post-translationally modified proteins, which are implicated in disease pathogenesis [2]. One such modification is citrullination, the post-translational conversion of arginine to citrulline, which sparks pathogenic anti-citrullinated protein autoantibody production by B cells and autoreactive T cell activation, which can sustain joint inflammation [1]. Another modification is carbamylation, the chemical transformation of lysine into homocitrulline [2]. Homocitrullinated proteins (HomoCitP) have been shown to elicit autoantibody responses highly specific to RA, to correlate with more severe disease phenotypes, and implicate B cells in the immune response to homocitrulline [3–5].

Given that B cells require cognate CD4+ T-cell help to produce high-affinity autoantibodies, the presence of IgG anti-homocitrullinated protein antibodies (AHCPA) in RA strongly implies that CD4+ T-cell responses to HomoCitP exist [3,6]. We previously used *in silico* modeling to demonstrate that HomoCitP are accommodated within the peptide-binding groove of major histocompatibility complex (MHC) class II molecules encoded by RA-associated human leukocyte antigen (HLA)-DRB1 alleles [5]. These alleles encode a conserved five-amino-acid sequence known as the “shared epitope” (SE), which is the strongest genetic risk factor for RA and is present in up to 60% of patients [1]. We have previously shown that splenocytes from mice expressing the SE proliferate in response to HomoCitP, supporting the hypothesis that SE-expressing MHC-II presents HomoCitP to CD4+ T helper (Th) cells [4]. However, the phenotype of these cells remains poorly understood.

Elevated levels of the CD4+ Th-cell subsets, Th1 and Th17, and their associated cytokines have been consistently observed in RA patients, suggesting their role in driving inflammation and joint destruction [1,7,8]. In addition, Th cells provide crucial “help” signals through dendritic cell licensing, co-stimulation, and cytokine production for the expansion and functional programming of CD8+ T cells [9–13]. This help enhances cytotoxic activity and supports the formation of robust memory responses, which are critical in sustaining chronic inflammation [13]. Recent single-cell whole transcriptome analyses revealed that clonally expanded CD8+ T cells, activated by citrullinated antigens, exhibit cytotoxic activity and are present in the RA synovium [14]. These findings highlight the potential contribution of antigen-specific CD8+ T cells to RA pathogenesis [14], prompting further investigation into whether similar responses occur in the context of homocitrullinated antigens.

Recently, Choudhury et al. demonstrated that CD4+ T cells from RA patients can proliferate in response to HomoCitP [15]. However, they did not investigate the phenotypic or functional characteristics of these cells, nor did they examine potential CD8+ T cell involvement. Moreover, their work focused on human peripheral blood (PB) samples and did not discuss *in vivo* mouse models for homocitrulline. In contrast, our study uses a humanized SE-expressing HLA-DRB1*04 (DR4)-transgenic (DR4tg) mouse model, which allows for an in-depth investigation of immune responses in a physiologically relevant, whole-organism context [4]. The aim of our study was to comprehensively characterize CD4+ and CD8+ T-cell responses to HomoCitP, including activation, cytokine production, proliferation, memory and effector phenotypes, using this DR4tg model. This approach provides critical insights into the cellular mechanisms through which homocitrullinated peptides can contribute to RA pathogenesis.

## Methods

### 2.1 DR4tg Mice

Humanized HLA-DRB1*04-transgenic (DR4tg) mice on a C57BL/6 background [4,16] were maintained under pathogen-free conditions. All procedures complied with the Canadian Council on Animal Care guidelines and were approved by Western University’s Animal Care and Use Committee (AUP 2022-150) .

### 2.2 Antigens and immunizations

A HomoCitJED peptide containing 9 homocitrulline residues and a control peptide, LysJED, where homocitrulline was replaced with lysine were synthesized by Creative Peptides (Shirley, NY, USA) [3,4,17].

For immunizations, male and female mice aged 8-12 weeks were injected subcutaneously in flanks (50 μl/flank) with a total of 100 μg of HomoCitJED dissolved in 100 μL of phosphate-buffered saline (PBS), containing Complete Freund’s Adjuvant (CFA) with *Mycobacterium tuberculosis* H37RA at a final concentration of 2 mg/mL. Control mice received CFA in PBS.

### 2.3 Cell preparation for cytofluorimetric analyses

Spleens and inguinal draining lymph nodes (dLNs) were harvested on day 10 post-immunization, prepared into single-cell suspensions, and stained with indicated fluorochrome-conjugated monoclonal antibodies. Cells were analysed on a BD LSR II or FACSymphony A1. Details on antibody panels, gating strategies, median fluorescence intensity calculations, and controls are in Supplementary Methods, **Figure S1-2**, **Tables S1-2**.

### 2.4 Proliferation and cytokine detection in culture supernatants

Splenocytes and dLN cells were stained with CellTrace™ Violet proliferation dye and cultured with HomoCitJED, LysJED, or medium alone, followed by flow cytometry. Cell proliferation was quantitated as a stimulation index [SI = (% cells proliferated in samples with peptide) / (% of cells proliferated in samples with medium alone)] (SI ≥ 2.0 deemed positive). For splenocytes, supernatants were assayed for various cytokines. Details of the methods are included in Supplementary Methods, **Table S1**, and **Figure S3**.

### 2.5 Intracellular cytokine staining (ICS)

The ICS protocol was adapted from Lovelace and Maecker [18]. Briefly, splenocytes/dLN cells were incubated with either HomoCitJED or medium alone. Cells were stained for surface markers, fixed/permeabilized, and labelled intracellularly for IL-17A, IFN-γ, TNF-α and IL-6. Cytokine-producing cells were normalized to the medium-alone condition and reported as the [% of cytokine+ T cells (HomoCitJED) / % of cytokine+ T cells (medium alone)]. Full experimental details are available in the Supplementary Methods, **Table S1**, and **Figure S4**.

### 2.7 Statistics

Two-tailed Mann-Whitney U tests compared HomoCitJED-immunized and control groups and, where indicated, peptide-stimulated versus control cultures. Box-and-whisker plots display all data points (median, interquartile range (IQR), minima, maxima). Analyses were performed in GraphPad Prism 9; p < 0.05 was considered significant.

## Results

### 3.1. Th-cell responses to HomoCitP are skewed toward Th1 and Th17 phenotypes

The following Th-cell subsets in dLNs were examined: T-bet+ Th1 cells, RORγt+ Th17 cells, GATA-3+ Th2 cells, and FoxP3+ Tregs. In HomoCitJED immunized mice, the proportion of Th1 cells relative to total CD4+ T cells was 17-fold higher compared to controls (5.4% vs. 0.2%; *p*=0.0022). Also, HomoCitJED-immunized mice had significantly higher Th17 cell frequencies (22.3%) when compared to control animals (3.3%; *p*=0.0043). There were no differences in the proportions of Th2 and Treg cells between the groups (**Figure 1A-B**).

**Figure 1.**
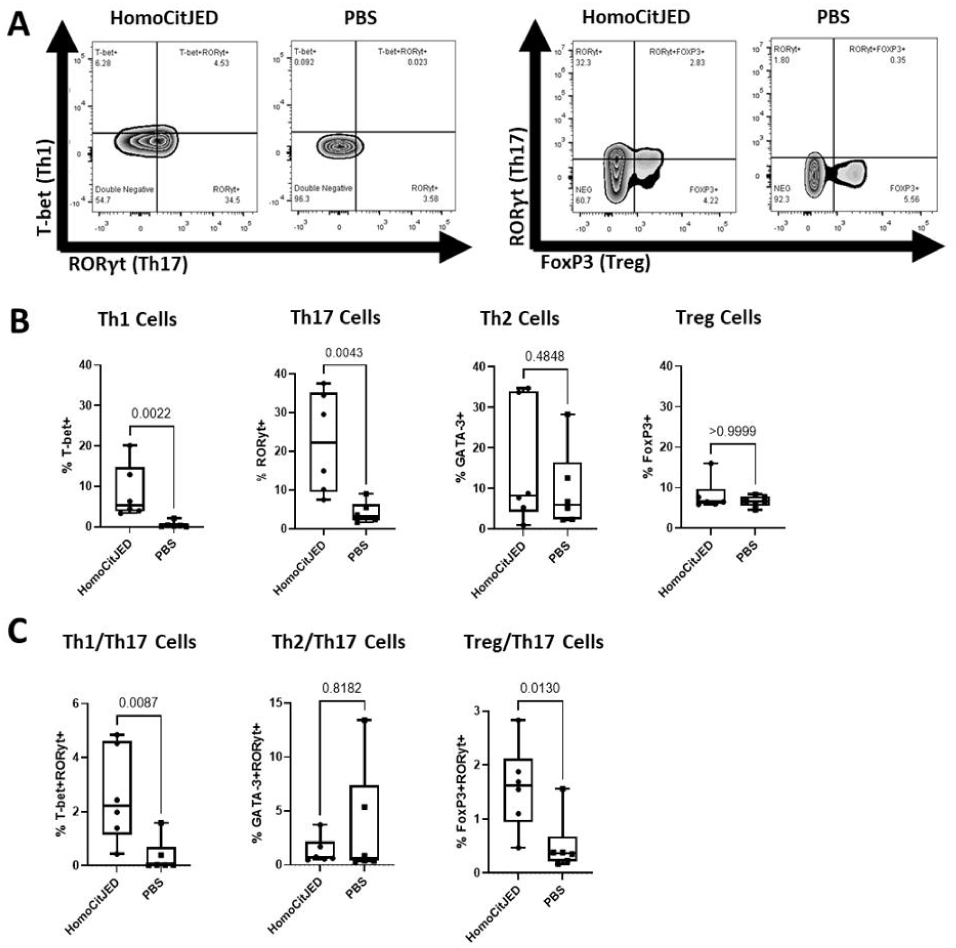
Th subsets in DR4tg mice after immunization with HomoCitJED. DR4tg mice were injected subcutaneously with CFA and either HomoCitJED or PBS. On day 10, dLNs were harvested, and their cells were analyzed by flow cytometry. Representative flow plots for Th1/Th17 and Treg/Th17 staining are shown in **A**. Quantification of T-bet+ Th1, RORγt+ Th17, GATA-3+ Th2, and FoxP3+ Treg cell frequencies is depicted in **B**. T-bet+RORγt+, GATA-3+RORγt+, and FoxP3+RORγt+ Th cells were also investigated and quantified in **C**. Values are % of total CD4+ T cells. Graphs show the median and IQR with each symbol representing an individual mouse (n=6/group). A p-value < 0.05 determined by the Mann-Whitney U test was considered significant.

Since Th17 cells can demonstrate plasticity, we also examined Th-cell subpopulations that simultaneously express the key transcription factors of more than one lineage. We found significantly higher proportions of T-bet+RORγt+ Th1/Th17 (2.2% vs. 0.02%; p=0.0087) and FoxP3+RORγt+ Treg/Th17 (1.6% vs. 0.4%; p=0.013) phenotypes in HomoCitJED-immunized mice compared to controls (**Figure 1A,C)**. No differences in GATA-3+RORγt+ Th2/Th17s were detected (**Figure 1C**).

### 3.2. HomoCitP immunization drives concurrent CD4+ T cell activation and exhaustion, signaling functional dysregulation

We evaluated the expression of T-cell activation markers (CD69 and CD25) in CD4+ T cells from dLNs. The frequency of CD4+ T cells expressing the early activation marker CD69 was comparable between the groups. In contrast, the proportion of CD4+ T cells positive for CD25, a later-appearing activation marker, was significantly higher in the HomoCitJED group (2.5 % vs 1.4 %; p = 0.0022) (**Figure 2A-B**).The proportions of activated CD25+ T-bet+ and RORγt+ T cells were significantly higher in HomoCitJED-immunized mice compared to controls (7.3% vs. 0.5%, p=0.0022 and 33.4% vs. 8.3%, p=0.0152, respectively) (**Figure 2C-D**). Conversely, the proportion of activated GATA3+ cells was significantly lower in mice injected with HomoCitP compared to controls (4.5% vs. 12.4%, p=0.0087) (**Figure 2D**). A significantly higher proportion of CD25+ T-bet/RORγt+ cells was observed in HomoCitJED-immunized mice (5.1%) compared to controls (0.1%, p=0.0260), whereas no significant difference was detected for CD25+ Th2/Th17 cells (2.5% vs. 1.3%, p=0.8182) (**Figure 2C-D**).

**Figure 2.**
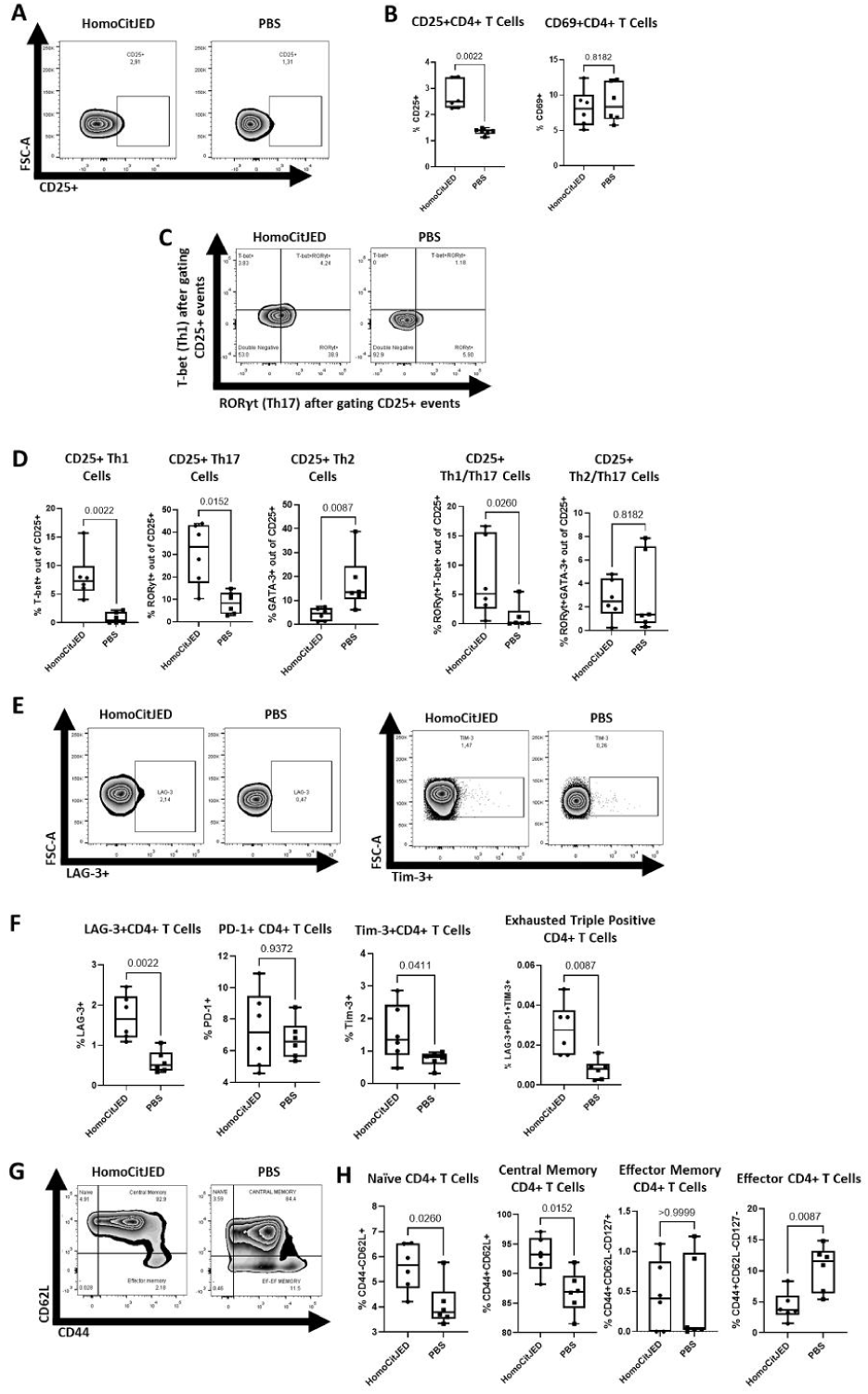
Activation, exhaustion, and memory phenotypes in DR4tg mice after immunization with HomoCitJED. DR4tg mice were injected subcutaneously with CFA and either HomoCitJED or PBS; cells were isolated from dLNs on day 10 and analyzed by flow cytometry. Representative flow plots of CD25+ staining are shown in **A**, and quantification of CD69+ and CD25+ CD4+ T cells is depicted in **B**. Representative plots of T-bet+, RORγt+, and T-bet+RORγt+ cells after CD25+ gating are shown in **C**. These subsets, along with GATA-3+ and GATA-3+RORγt+ Th cells, were quantified after gating on CD25+ and shown in panel **D**. FoxP3+ cells were excluded from analyses in A-D. Representative flow plots for LAG-3+ and Tim-3+ CD4+ T cells are shown in **E**, and their quantification is presented in **F**. Representative CD44/CD62L flow plots demonstrate an increase in central memory CD4+ cells in dLNs cells in **G**. Quantification of CD44+CD62L+ central memory (Tcm), CD44+CD62L-CD127+ effector memory (Tem), and CD44+CD62L-CD127-effector (Teff) CD4+ is presented in **H**. Values represent percentages of total CD4+ T cells or percentages of CD25+ Th cells for D. Graphs show the median and IQR; each symbol represents an individual mouse (n=6/group). A p-value < 0.05 by the Mann-Whitney U test was considered statistically significant.

Prolonged antigen exposure and chronic inflammation can induce T-cell exhaustion [19], which was also examined in this study. The proportions of lymphocyte activation gene-3 (LAG-3)+ and T-cell immunoglobulin and mucin-domain-containing protein 3 (Tim-3)+ CD4+ T cells in HomoCitJED-immunized mice were higher than in controls (1.7% vs. 0.5%; p=0.0022 and 1.4% vs. 0.8%, p=0.0411, respectively), whereas programmed cell death protein 1 (PD-1)+ CD4+ T cells were not significantly different between groups (**Figure 2E-F**). The proportion of CD4+ T cells that expressed LAG-3+ PD-1+ Tim-3+ (terminally exhausted [20]) was very low - 0.03% in HomoCitJED-immunized mice and 0.008% in controls (**Figure 2F**). Lastly, LAG-3 surface expression was significantly upregulated on CD4+ T cells from HomoCitJED-immunized mice compared to controls(MFI of 473 vs. 240, p=0.0087) (**Figure S5**).

We also investigated memory phenotypes and identified central memory T cells (Tcm) as CD44+CD62L+, effector memory T cells (Tem) as CD44+CD62L-CD127+ cells, terminal effector T cells (Teff) as CD44+CD62L-CD127-cells, and naïve T cells as CD44-CD62L+ [21,22,23]. The proportion of CD4+ Tcm cells was 93.2% in HomoCitJED-immunized mice compared to 86.9% in controls (p = 0.0152) (**Figure 2G-H**). For Tem cells, the proportion of these cells was low overall (0.4%) and there were no significant differences between the groups. As for Teff cells, these were significantly lower in HomoCitP mice (3.7%) compared to controls(11.6%; p=0.0087) (**Figure 2G-H**). Lastly, mice immunized with HomoCitJED had a higher proportion of naïve CD4+ T cells (5.7%) compared to controls (3.8%; p = 0.0260) (**Figure 2G-H**).

### 3.3 Immunization with HomoCitP promotes CD8+ T-cell activation and exhaustion

Increasing evidence suggests that CD8+ T cells play a role in RA [14]. Therefore, we investigated activation, exhaustion, and memory phenotypes, as well as effector functions of CD8+ cells in response to HomoCitP in dLNs of DR4tg mice.

With respect to early activation, there were no differences in CD69-expressing CD8+ T-cell frequencies. However, there was a higher proportion of CD25+CD8+ T cells in mice immunized with HomoCitJED (9.5%) compared to controls (1.6%; p=0.0037) (**Figure 3A-B**). There were also higher proportions of CD8+ T cells expressing LAG-3 (19% vs. 4.7%, p=0.0059), PD-1 (9.5% vs. 5.8%, p=0.0382), and Tim-3 (3.1% vs. 0.6%, p=0.0205), indicating an increase in exhaustion phenotypes (**Figure 3C-D**). Similar to CD4+ cells, the co-expression of all three markers (terminally exhausted cells) was very low: 0.08% in HomoCitJED-immunized mice and 0.02% in controls. In addition, CD8+ T cells from HomoCitJED-immunized mice displayed elevated Tim-3 surface expression relative to controls (MFI of 2357 vs. 2299, p = 0.0379) (**Figure S6**).With respect to memory and effector CD8+ cells, Tem cells (12.5%) were higher in HomoCitJED vs. controls (7.5%); p=0.0200, whereas Tcm and Teff cells were not different (**Figure 3E-F**).

**Figure 3.**
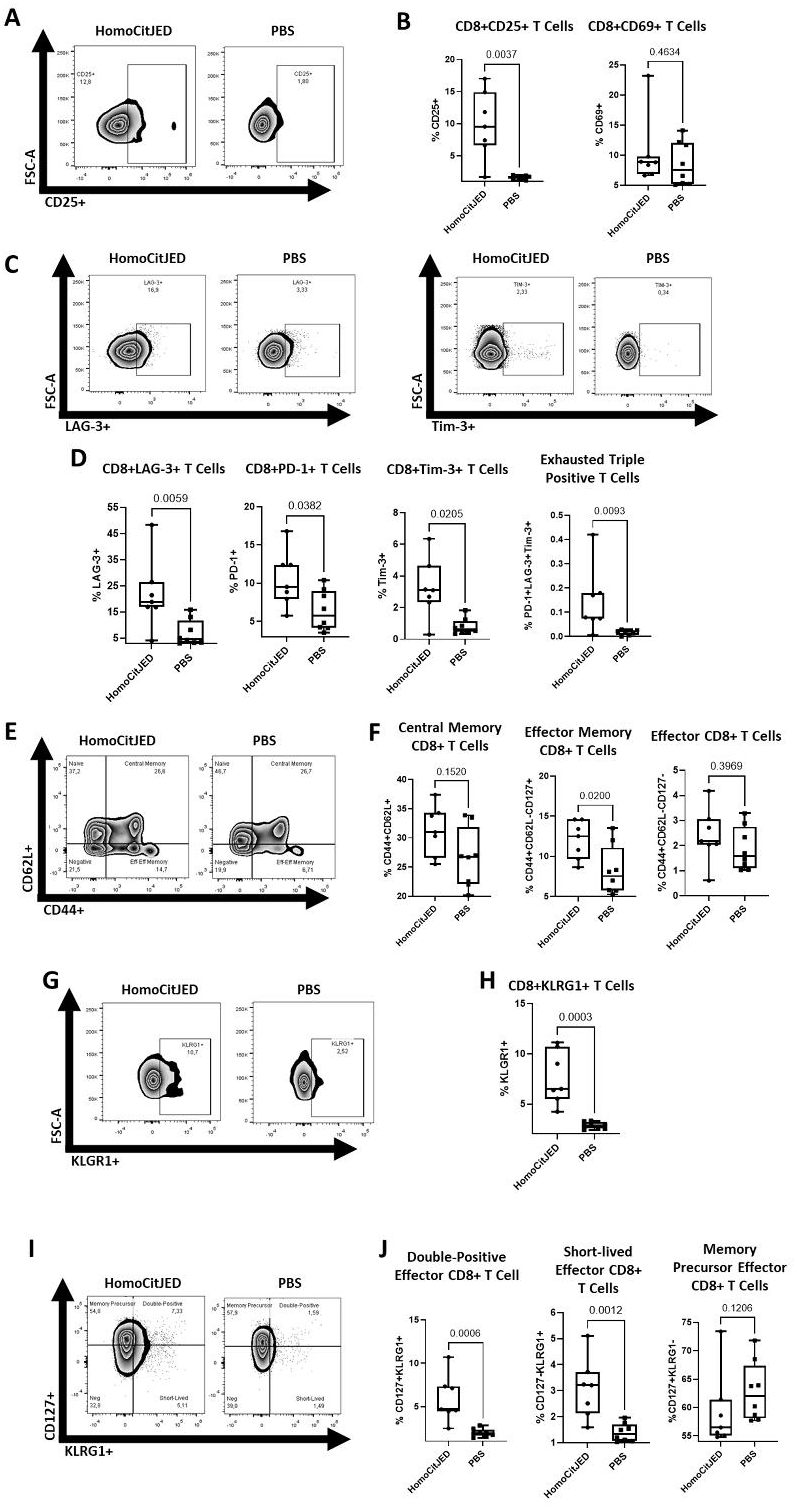
CD8+ T-cell phenotypes in DR4tg mice after immunization with HomoCitJED. DR4tg mice were subcutaneously immunized with CFA and either HomoCitJED or PBS, and cells were isolated from the dLNs on day 10 followed by flow cytometry analysis. Representative staining for CD25+ CD8+ T cells is shown in **A**, and quantification of activation markers CD69+ and CD25+ CD8+ T cells is depicted in **B**. Representative flow plots for LAG-3+ and Tim-3+ expression are shown in **C**, while quantification of LAG-3+, PD-1+, Tim-3+, and triple-positive exhausted cells is shown in **D**. Representative flow plots for CD44/CD62L expression are shown in **E**. CD44+CD62L+ Tcm, CD44+CD62L-CD127+ Tem, and CD44+CD62L-CD127-Teff CD8+ T cells were quantified in **F**. Increased KLRG1 expression is shown in **G**, and the corresponding quantification is shown in **H**. KLRG1 and CD127 surface markers were used to define DPECs (CD127+KLRG1+), SLECs (CD127-KLRG1+), and memory precursor T cells (CD127+KLRG1-) in **I**, which were quantified in **J**. Values represent the percentage of total CD8+ T cells. Graphs display the median and IQR; each symbol represents an individual mouse (n=7-8/group). A p-value < 0.05 by the Mann-Whitney U test was considered statistically significant.

We next examined CD8+ T cells expressing killer cell lectin-like receptor G1 (KLRG1). Although KLRG1 often functions as an inhibitory receptor, it is also expressed on CD8+ T cells that produce inflammatory mediators in autoimmune responses [25]. HomoCitJED immunization significantly increased the frequency of KLRG1+ CD8+ T cells compared with controls (6.5% vs. 2.8%; p = 0.0003) (**Figure 3G–H**). To further characterize these cells, we assessed their co-expression with the IL-7 receptor α-chain (CD127). T cells that are KLRG1+ CD127+ are referred to as double-positive effector cells (DPECs); they have limited capacity to become long-lived memory cells [26,27]. KLRG1+ CD127-cells, known as short-lived effector cells (SLECs), lack CD127, which is required for long-term survival. SLECs are specialized for immediate effector function [26,27]. In contrast, KLRG1-CD127+ cells are memory precursor T cells with the potential to differentiate into Tcm and Tem’s [26,27].

We investigated these phenotypes and observed that, compared with controls, HomoCitJED immunization led to significantly higher proportions of SLECs (3.2% vs. 1.3%; p = 0.0012) and DPECs (4.7% vs. 1.9%; p = 0.0006). The frequency of memory precursor T cells, however, remained unchanged (**Figure 3I–J**).

### 3.4. CD4+ and CD8+ T cells mount detectable recall proliferation to HomoCitJED

In RA, specific autoantigens, including HomoCitP, are recognized by T cells, leading to their activation and proliferation [15,28]. However, existing studies are limited and often do not investigate CD8+ T cells [15,28]. To assess whether HomoCitP can stimulate both CD4+ and CD8+ T-cell proliferation, we evaluated the recall responses of these subsets *ex vivo*.

HomoCitJED-immunized mice (5/5, 100 %) showed a positive proliferative response, whereas none of the controls did (0/5); median CD4+ stimulation indices were 2.7 versus 1.1, respectively (p = 0.0079; **Figure 4A-B**). For CD8+ cells, 5/5 (100%) HomoCitJED-immunized mice exhibited responses to homocitrulline, compared to 0/5 (0%) in controls, with SI 3.8 vs. 0.6 (p=0.0079), respectively (**Figure 4C-D**).

**Figure 4.**
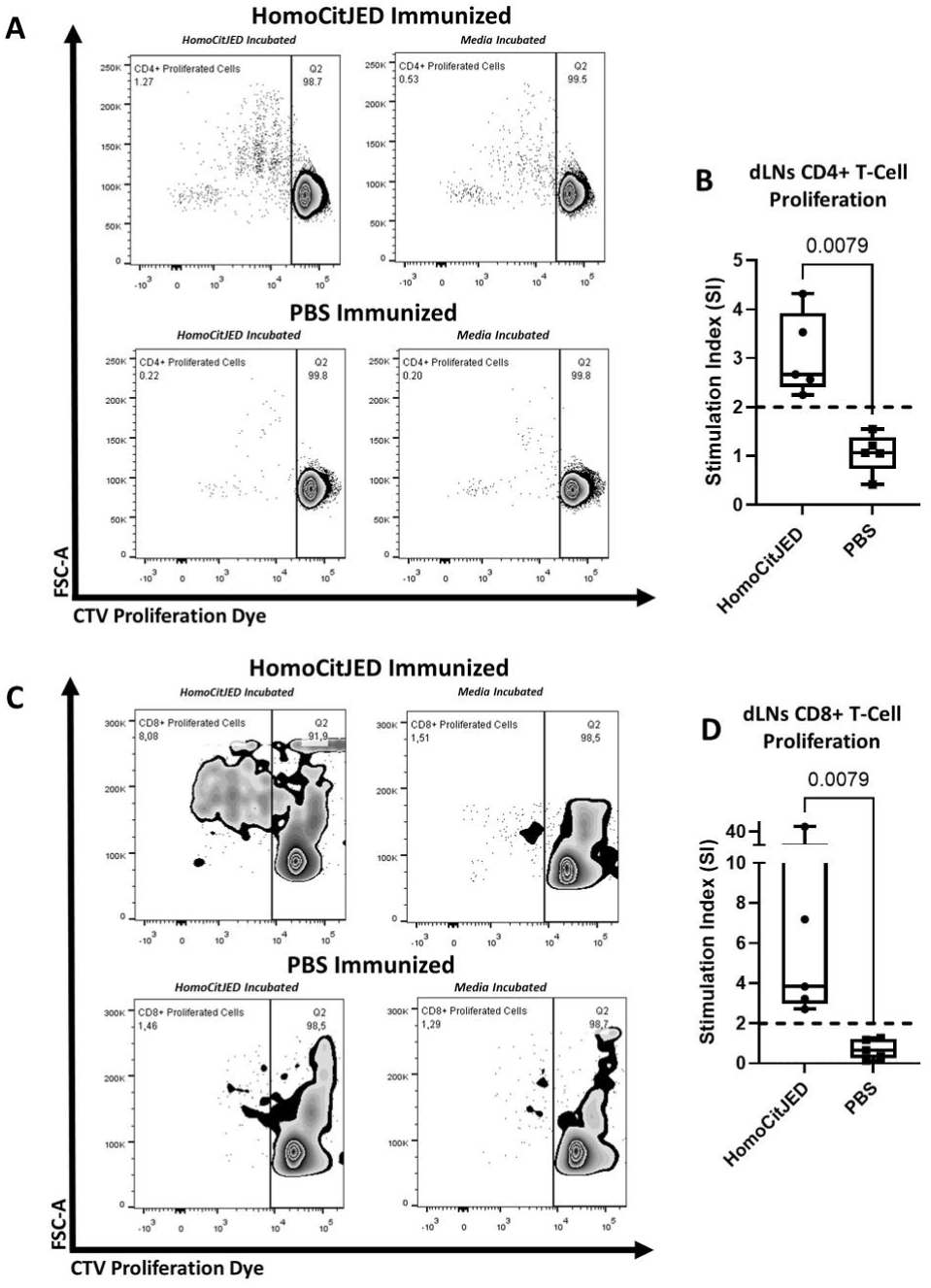
Cognate T-cell proliferation in HomoCitJED-immunized mice. dLN cells from HomoCitJED-injected and control mice were incubated with 100 μg/mL HomoCitJED or medium alone and analyzed by flow cytometry 72 hours later. Representative flow plots of CD4+ T-cell proliferation in each group are shown in **A**, with corresponding stimulation indices (SI) summarized in **B**. Representative plots for CD8+ T-cell proliferation are shown in **C**, with SI data presented in **D**. SI was calculated as the percentage of proliferating cells in peptide-stimulated samples divided by that in medium-only samples. A SI > 2.0 (indicated by a dashed line) was considered a positive proliferative response. Each symbol represents pooled cells isolated from the dLNs of 3 mice (n=15/group). Graphs show the median and IQR. Statistical significance was determined by the Mann-Whitney U test; p < 0.05 was considered significant.

Additionally, we tested for proliferative responses to the backbone of HomoCitJED using a control peptide, LysJED, which yielded no proliferation in HomoCitJED-immunized mice, confirming that the observed proliferation was specific to homocitrulline (**Figure S8**).

### 3.5. HomoCitJED immunization enriches IL-17A-producing Th1/Th17 cells in draining lymph nodes

Cytokines regulate a wide range of inflammatory processes involved in the pathogenesis of RA [29]. Therefore, we examined the cytokines produced by T cells from dLNs stimulated *ex vivo* with HomoCitJED. Analysis of single-cytokine producers showed that HomoCitJED-immunized mice had a significantly higher ratio of IL-17A+ CD4+ T cells than controls (median normalized to media = 2.20 vs 1.16; p = 0.007) (**Figure 5A**). In contrast, TNF-α+, IFN-γ+ and IL-6+ CD4+ T cells did not differ between the groups (**Figure 5A**).

**Figure 5.**
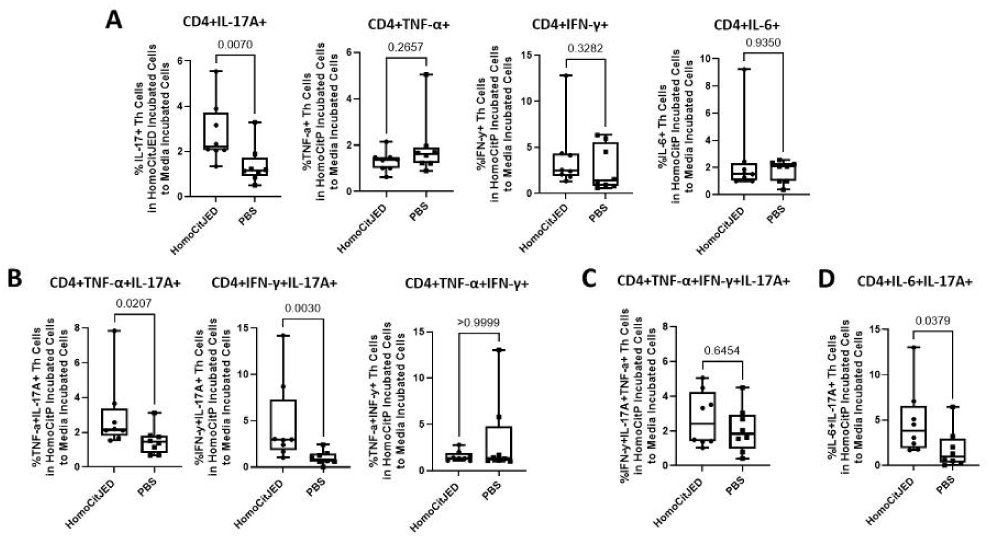
Cytokine profiles of CD4+ T cells from dLNs after ex vivo stimulation with HomoCitJED. DR4tg mice were immunized with HomoCitJED or PBS; dLN cells were harvested on day 10 post-immunization and cultured with 100 µg/mL HomoCitJED (or medium alone) for 5 h. Intracellular staining for TNF-α, IFN-γ, IL-6 and IL-17A was analyzed by flow cytometry. Cells producing a single cytokine are shown in **A**. Cells producing Th1-type cytokines or both a Th1 and a Th17 cytokine are represented in **B**. Cells producing three cytokines characteristic of Th1/Th17 responses are shown in **C**. Cells co-expressing IL-17A with IL-6 are shown in **D**. Cytokine production is expressed as the ratio of cells stimulated with HomoCitJED to those cultured in medium alone. Graphs display the median and IQR; each symbol represents an individual mouse (n=8/group). Statistical significance was determined by the Mann-Whitney U test; p < 0.05 was considered significant.

When we evaluated dual-cytokine producers, HomoCitJED-immunized mice showed significantly higher ratios of TNF-α+IL-17A+ and IFN-γ+IL-17A+ cells than controls (2.15 vs 1.44, p = 0.0207; and 2.94 vs 0.81, p = 0.003, respectively) (**Figure 5B**). Triple-positive (TNF-α+IFN-γ+IL-17A+) CD4+ T cells were comparable between the groups (**Figure 5C**). Finally, the proportion of IL-6+IL-17A+ CD4+ T cells was significantly higher in HomoCitJED-immunized mice (3.79 vs 0.94; p = 0.0379) (**Figure 5D**).

To determine whether CD8+ T-cell (Tc) subsets showed a similar cytokine profile, we performed the same analyses on these cells. In the single cytokine producers, HomoCitJED immunization increased the ratio of only IL-17A+ CD8+ T cells compared with controls (2.24 vs 0.62; p = 0.0104) (**Figure 6A**). A significant enrichment of IFN-γ+IL-17A+ CD8+ T cells, characteristic of Tc1/Tc17 cells was observed in HomoCitJED-immunized mice (2.46 vs 0.78; p = 0.0368), whereas TNF-α+IFN-γ +, TNF-α+IL-17A+ and IL-6+IL-17A+ subsets were comparable between groups (**Figure 6B**). TNF-α+IFN-γ+IL-17A+ CD8+ T cells also did not differ between HomoCitJED and controls (**Figure 6C**).

**Figure 6.**
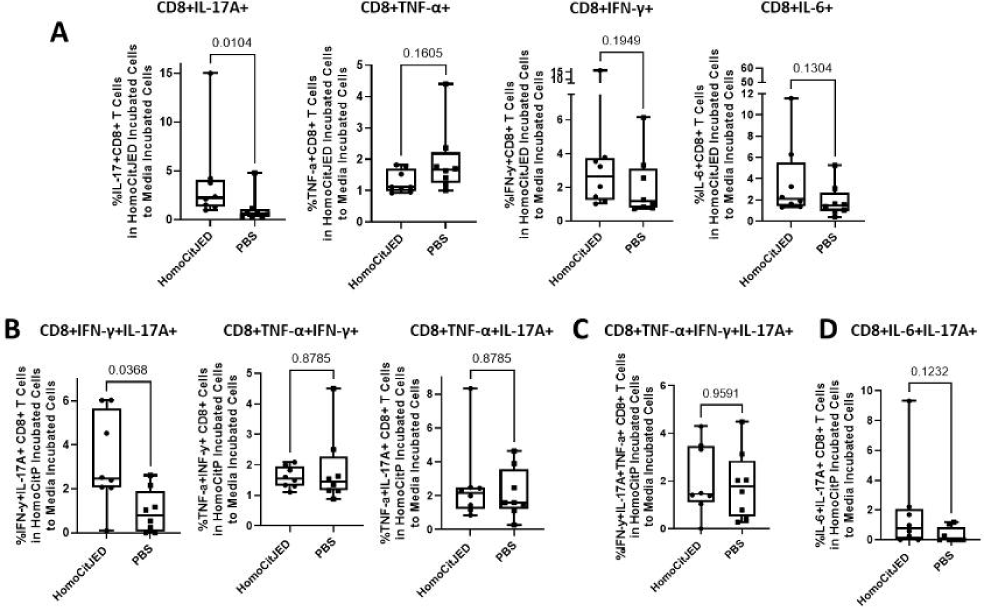
Cytokine profiles of CD8+T cells from dLNs after ex vivo stimulation with HomoCitJED. DR4tg mice were immunized with HomoCitJED or PBS; dLN cells were harvested on day 10 post-immunization and cultured with 100 µg/mL HomoCitJED (or medium alone) for 5 h. Intracellular staining for TNF-α, IFN-γ, IL-6 and IL-17A was analyzed by flow cytometry. Cells producing a single cytokine are shown in **A**. Cells producing cytotoxic T-cell type 1 (Tc1) cytokines or both Tc1 and Tc17 cytokines are denoted in **B**. Cells producing three cytokines associated with Tc1/Tc17 responses are represented in **C**. Cells co-expressing IL-17A with IL-6 are shown in **D**. Cytokine production is expressed as the ratio of cells stimulated with HomoCitJED to those cultured in medium alone. Graphs display the median and IQR; each symbol represents an individual mouse (n=8/group). Statistical significance was determined by the Mann-Whitney U test; p < 0.05 was considered significant.

### 3.6. Splenic T-cell responses in HomoCitJED-immunized mice are pro-inflammatory for CD4+ T cells but minimal in CD8+ T cells

To assess not only local responses in dLNs but also systemic responses, we analyzed splenic T cells. HomoCitJED injection enriched FoxP3+RORγt+ and CD25+CD4+ T cells compared with controls, while no phenotypic differences were observed in CD8+ T cells. However, the peptide induced homocitrulline-specific proliferation of both CD4+ and CD8+ splenic T cells. In addition, splenocytes from HomoCitJED-injected mice released significantly higher levels of pro-inflammatory cytokines (IL-17A, TNF-α, IFN-γ, and IL-2) compared with controls. By contrast, IL-10 concentrations did not differ between groups. Finally, a higher proportion of CD4+ T cells producing IL-6, IL-17A, and IFN-γ/IL-17A was detected in HomoCitJED-injected mice, whereas no differences were observed in CD8+ T cells. Full data are provided in the **Supplementary Results**.

## Discussion

T cells play a pivotal role in the pathogenesis of RA [8]. Although T-cell reactivity to HomoCitP has received less attention, antibodies targeting HomoCitP are readily detectable in RA patients years before clinical onset, and their presence predicts more erosive disease [5,30,31]. Mass-spectrometry studies have also mapped multiple homocitrullinated epitopes in RA joints, potential targets for RA-specific T and B-cell responses [32]. Our study revealed that immunization with HomoCitP triggered a robust, pro-inflammatory CD4+ T-cell response, reshaped CD8+ T-cell phenotypes, and drove clonal T-cell expansion in HLA-DR4tg mice, a humanized animal model of RA, supporting a potential pathogenic role in human disease.

After injection with HomoCitJED, mice exhibited significant skewing toward Th1 and Th17 responses, consistent with RA as a Th1- and Th17-driven disease [1,8]. CD4+ T cells in DR4tg mice preferentially produced IFN-γ and TNF-α, known to sustain Th1 inflammatory responses that are predictive for RA onset [1,8], in concert with IL-17A. This cytokine signature is characteristic of hybrid Th1/Th17 cells, reflecting the plasticity in which Th17 cells progressively acquire Th1 traits [1]. This is also supported by our phenotyping data, which showed a marked expansion of T-bet+RORγt+ Th1/Th17 cells. Thus, together our data suggest that homocitrullinated antigens propel CD4+ T cells toward a Th1/Th17 hybrid phenotype, amplifying both IL-17A- and IFN-γ/TNF-α-mediated inflammatory pathways rather than generating separate, classical Th1 effectors. *In vitro* studies by others have shown that Th17 cells can re-programme into more pathogenic “Th17/1” cells, characterized by both RORγt/IL-17 and T-bet/IFN-γ production and an increased proliferative capacity [1,33]. The observed plasticity is largely unidirectional, as committed Th1 cells rarely revert to a Th17 phenotype [33,34]. Clinically, circulating Th1/Th17 cells were found to be elevated in RA patients relative to healthy controls, underscoring their relevance to disease onset [35, 36]. The specific antigens driving these Th1/Th17 cells in RA were previously unknown, and our data point to HomoCitP as viable candidates.

Th17/Treg cells are another Th subset identified in RA patients but not extensively researched [1]. These cells can exhibit pro-inflammatory (Th17-like) or anti-inflammatory (Treg-like) properties [1,33]. Zhang et al. found elevated Th17/Treg cells in RA patients’ PB compared to controls, lacking suppressive function in synovial fluid (SF), indicating a pro-inflammatory phenotype [37]. We observed an expansion of Th17/Treg cells in HomoCitJED-immunized mice, along with higher IL-17A, whereas Treg frequency and IL-10 production in supernatants were unchanged. Together, these findings suggest that HomoCitP tilts the Th17/Treg balance toward a pro-inflammatory, Th17-dominant phenotype that could further fuel RA pathology.

Whereas many of our observations echo established RA immunopathology [8], one feature appears distinctive for the HomoCitP response: the Treg compartment appeared to be preserved. Specifically, IL-10 concentrations in the supernatants after exposure to HomoCitJED remained low and unchanged. Thus, the strong Th1/Th17 skew is not accompanied by the marked Treg contraction or compensatory IL-10 surge often reported in RA [39]. This preservation of Tregs may be unique to the HomoCitP response, reflecting a predominantly pro-inflammatory reaction that fails to engage regulatory pathways.

We also observed a marked expansion of activated CD25+ CD4+ T cells in both dLNs and spleens in response to HomoCitP. This activation signature was reinforced by two functional readouts: the supernatants from splenocytes treated *ex vivo* with HomoCitJED contained significantly higher IL-2, the growth factor signaling through CD25, and CD4+ T cells showed enhanced proliferation in response to HomoCitJED. Patients with RA are known to harbour activated T cells in both PB and SF, and these cells proliferate in response to citrullinated antigens [15,28]. Our data extend this finding by showing that HomoCitP likewise drive strong T-cell activation and proliferation. Further analysis showed that within the activated (CD25+) CD4+ population, Th17, Th1 and Th1/Th17 cells were higher, whereas Th2 and Th2/Th17 frequencies were lower in HomoCitJED-immunized mice. The observed imbalance among Th2, Th1 and Th17 immunity has long been implicated in RA pathogenesis [8,38], and our data indicate that exposure to HomoCitP can contribute to this imbalance.

HomoCitJED also activated CD8+ T cells. We detected a significant increase in CD25-expressing CD8+ T cells in both dLNs and spleens, mirroring the elevated CD8+CD25+ population reported in RA patients’ PB where its frequency was shown to correlate with clinical severity [40]. These findings extend our earlier work by highlighting that HomoCitP trigger both T-cell lineages, broadening the pool of potentially pathogenic effectors in RA [4].

Persistent antigen exposure usually pushes T cells into exhaustion, diminished proliferation and cytokine secretion coupled with heightened expression of inhibitory receptors. Strikingly, HomoCitP did not follow that paradigm in our DR4tg model. Both CD4+ and CD8+ compartments up-regulated exhaustion markers, such as Tim-3 and LAG-3; yet, instead of functional loss, we observed IL-17A/TNF-α/IFN-γ production, proliferation, and a shift toward effector-memory phenotypes. This combination of exhaustion markers in hyper-active cells reveals a dysregulated checkpoint landscape that may allow inflammation to persist unchecked. Luo *et al.* previously reported an increase in exhausted Th cells in RA patients, and Ponchel *et al.* described hyper-responsive T cells prone to excessive cytokine release and proliferation [41,42]. Our data build upon those clinical observations by supporting HomoCitP as a potential trigger simultaneously driving activation and subverting normal exhaustion programmes, an imbalance that may contribute to the pathogenesis of RA.

Given the role of memory and effector T cells in chronic disease, we investigated the response of these cells to HomoCitP [8,12,13]. We observed an increase in CD4+ Tcm cells and a decrease in CD4+ Teff cells, alongside an increase in CD8+ Tem cells in dLNs after HomoCitJED immunization. This suggests different memory components are activated for CD4+ and CD8+ T cells at this early time point. Specifically, long-lasting CD4+ Tcm cells, which remain in lymphoid organs, may supply new T cells responding to homocitrulline, contributing to chronic disease. Effector cells are short-lived, and migrate from lymphoid tissues to peripheral sites, complicating single-time-point assessments. The observed decrease in Teff cells may reflect their migration to peripheral sites where they interact with targets. Our findings align with the observation of James et al. showing higher citrulline-specific memory T cells in RA patients and Cho et al. who reported increased CD8+ Tem cells in SF [7,43].

We also investigated KLRG1, an immune checkpoint marker. Although KLRG1 often functions as an inhibitory receptor that raises the activation threshold and can limit autoimmunity, it is also expressed on T cells that produce inflammatory mediators in autoimmune responses [25]. Altered KLRG1 expression has been reported in several autoimmune conditions, including RA where KLRG1+ T cells positively correlate with disease severity [25] and poor drug response [44]. Our findings show increased KLRG1+ CD8+ (SLECs and DPECs) T cells in HomoCitJED-injected mice. SLECs have immediate effector function but short lifespans and low proliferative potential. They contribute to inflammation in autoimmune diseases like type 1 diabetes, and may be linked to RA autoimmunity [25,45]. DPECs appear to positively impact immune response in follicular lymphoma and may have a role in driving inflammatory responses [27]. Overall, these changes in CD8+ cells uncover phenotypes that may be involved in the pathophysiological changes in RA. Further research is needed to fully understand these cells.

In our HLA-DR4 humanized mouse model, animals express RA-associated MHC II molecules, which suggests that antigen presentation first activates CD4+ T cells. These Th cells can then stimulate CD8+ T-cell maturation [9–13,46], suggesting that changes in CD8+ T-cell phenotypes are driven by DR4-specific CD4+ T-cell activation. This is supported by CD8+ T-cell expression of KLRG1, CD25, and CD44 (markers expressed after CD4 help) [12, 48]. Another alternative is direct presentation of the 13-mer HomoCitJED peptide on MHC I molecules, which is less likely given that MHC I usually binds 7-11-mer ligands. Determining whether CD8+ cells rely solely on DR4-restricted CD4+ help, or can also recognize homocitrulline directly, remains essential for clarifying RA pathogenesis and guiding targeted therapies.

Our study has several strengths. We utilized the peptide HomoCitJED, which captures reactivity and generates immune responses toward various post-translationally modified antigens [3,5]. This approach is reflective of individuals with RA, as reactivity toward different post-translationally modified antigens has been observed in this population [3,5]. Additionally, the DR4-transgenic mouse model employed in this study is highly relevant to human RA, as it bears the strongest genetic risk factor for the disease [1]. Our team previously detected HomoCitJED immune responses in these mice and showed that they develop RA-like arthritis at later time points [4,49].

While our findings constitute a significant step toward defining homocitrulline-driven immunity in RA, there are several limitations. First, because our analyses stop on day 10 post-immunization, longitudinal studies are needed to follow HomoCitJED-specific T- and B-cell responses (and T/B-cell interactions) through to the onset and progression of arthritis. Second, we have yet to test causality: adoptive-transfer or subset depletion approaches will be required to determine whether the populations we identified are necessary, or merely permissive, for disease progression. By outlining these directions, our study not only delivers the first integrated view of CD4+ and CD8+ responses to homocitrullinated antigen in a genetically relevant model but also lays the groundwork for future studies clarifying how homocitrulline-driven immunity shapes disease timing, severity and, ultimately, therapeutic targeting.

In summary, our findings support the ability of SE-expressing MHC Class II molecules to present homocitrullinated antigens, promoting pro-inflammatory T-cell phenotypes linked to RA pathogenesis. Additionally, we highlight both CD4+ and CD8+ T-cell involvement, showing activation and exhaustion in response to HomoCitJED, suggesting a dual role in inflammation and immune regulation. Overall, our study underscores the role of HomoCitP in shaping T-cell responses in RA and provides insights for potential T-cell-targeted therapies.

## Acknowledgements and affiliations

We extend our sincere gratitude to Garth Blackler for his assistance with animal work training. This work was previously presented at the 2025 Canadian Rheumatology Association Annual Scientific Meeting (Ulanova S, Buckley G, Haeryfar SM, Cairns E, Barra L. Investigating Homocitrulline-Specific T-Cell Responses in Rheumatoid Arthritis. J Rheumatol. 2025;52(Suppl 2):37–38. doi:10.3899/jrheum.2025-0314.TOUR7C) and at the 19th International Congress of Immunology, August 2025 (Frontiers in Immunology, accepted).

## Declaration of generative AI and AI-assisted technologies in the manuscript preparation process

During the preparation of this work the author(s) used Microsoft Copilot/ChatGPT in order to help improve language and readability of the manuscript. After using this tool/service, the author(s) reviewed and edited the content as needed and take(s) full responsibility for the content of the published article.

