## Supplementary Materials for "Homocitrullinated Peptides Drive Pro-Inflammatory T-Cell Responses in a Humanized Mouse Model of Rheumatoid Arthritis"

### **Supplementary Methods**

### **1.1 Mice and Antigens**

Humanized HLA-DRB1*04:01-transgenic (DR4tg) mice on a C57BL/6 background [1,2] were bred in-house. Mice were housed in specific pathogen-free conditions at the Animal Care and Veterinary Services facility at the University of Western Ontario under controlled temperature (20–22°C) and a 12-hour light/dark cycle, with ad libitum access to food and water.

All peptides used in the study were synthesized by Creative Peptides (Shirley, NY, USA). Peptide purity was assessed by high-performance liquid chromatography (HPLC), and sequence identity was confirmed by mass spectrometry. The supplier reported a purity of 94.5% for HomoCitJED and 92.08% for LysJED.

### **1.2 Cell preparation for cytofluorimetric analyses**

Mice were euthanized on day 10 post-subcutaneous (s.c.) injection with CFA and either HomoCitJED or PBS, and spleens along with inguinal draining lymph nodes (dLNs) were harvested into cold complete Roswell Park Memorial Institute 1640 medium (cRPMI), supplemented with 10% heat-inactivated fetal bovine serum (FBS), 1% Glutamax, 1% HEPES, 1% non-essential amino acids, 1% sodium pyruvate, and 1% penicillin/streptomycin (all from ThermoFisher). Splenocytes were obtained by manual disruption using a Wheaton Dounce tissue grinder. dLNs were processed into single-cell suspensions by gently pressing them through a 70-μm cell strainer with the plunger of a sterile syringe. Red blood cell lysis was performed using freshly prepared ammonium-chloride-potassium (ACK) buffer, pH 7.2-7.4. Following isolation, 1 × 10⁶ cells from each mouse were transferred into FACS tubes, washed with cell staining buffer (BioLegend), and incubated with 10 μL of anti-mouse Fc receptor blocking reagent (Miltenyi Biotec) for 10 minutes at 4°C to reduce non-specific binding. Cell viability was assessed using a 1:1000 dilution of Fixable Viability Dye (BioLegend), and surface marker staining was carried out in staining buffer for 30 minutes at 4°C. For intracellular staining, the Intracellular Fixation and Permeabilization Buffer Set (ThermoFisher) was used according to the manufacturer's instructions.

All fluorochrome-conjugated monoclonal antibodies (mAb) used for flow cytometry are listed in **Table S1**. Gating strategies for identifying CD4+ and CD8+ T cells and their subsets are detailed in **Table S2** and **Figures S1–2**. Since CD25 is also constitutively expressed by Treg cells, we excluded FoxP3+ Th cells from the analysis of activated CD25+ T cells (**Figure S1**). Gates were determined using Fluorescence Minus One (FMO) controls. Data were collected using a BD LSR II flow cytometer at the London Regional Flow Cytometry Facility and analyzed with FlowJo software (Tree Star). Median fluorescence intensity (MFI) was calculated using FlowJo software by gating on the specified cell population and extracting the median value of the fluorescence signal. The values were used to determine relative expression levels, with FMO controls applied to define positive populations.

### **1.3** **Splenocyte and dLN cell proliferation**

Cell suspensions of mouse spleens and dLNs, harvested on day 10 after s.c. injection with CFA and either HomoCitJED or PBS were stained with proliferation dye CellTrace™ Violet according to manufacturer’s instructions (ThermoFisher, **Table S1**). Cells were then cultured at 37°C at 5 × 10^5^ cells/well in media containing 100 μg/mL of HomoCitJED, control peptide LysJED, or cRPMI 1640 medium alone plus 0.1 mM of 2-mercaptoethanol (Gibco). Mitogens [10 ng/mL of Phorbol 12-myristate 13-acetate (PMA) plus 100 ng/mL of ionomycin (Abcam and Sigma)] were used as a positive control. After 72 hours of incubation, cells were stained with a live/dead stain, as well as anti-CD3, -CD4, and -CD8 antibodies and acquired on LSR II flow cytometer (**Table S1**). The gating strategy is outlined in **Figure S3**. Proliferative responses are reported as a Stimulation Index [SI = (% cells proliferated in samples with peptide) / (% of cells proliferated in samples with media alone)]. A cut-off value of 2.0 was considered indicative of a positive proliferative response. This threshold was determined based on the average percentage of proliferating cells in media alone plus two standard deviations (N=38 mice).

### **1.4 Quantification of cytokines in culture supernatants**

After a 48-hour incubation of splenocytes in proliferation conditions, 150 µl of supernatants were centrifuged at 1500 rpm for 10 minutes at 4°C, aliquoted, and stored at -80°C. Samples were later thawed and immediately used for cytokine quantification. IL-17A, IL-2, TNF-α, IFN-γ, and IL-6 levels were measured using ProQuantum™ high-sensitivity immunoassays (ThermoFisher), following the manufacturer’s instructions. Due to the unavailability of the ProQuantum kit for IL-10, the concentration of IL-10 was determined using a mouse IL-10 ELISA kit (ThermoFisher), following the manufacturer’s instructions. Results are presented as the concentrations of cytokines produced in HomoCitJED stimulation cultures normalized by subtracting the concentrations observed in the medium alone condition.

### **1.5 Intracellular cytokine staining (ICS)**

Ten days post-s.c. injection with CFA and either HomoCitJED or PBS, splenocytes and dLN cells from the mice were cultured at 37°C (2×10⁶ cells/well) for 5 hours with 100 μg/mL of HomoCitJED, medium alone, or PMA (10 ng/mL) + ionomycin (100 ng/mL) (positive control). A protein transport blocker, brefeldin A (eBiosciences), was added 2 hours after the start of incubation. Cells were then labelled for cell-surface and intracellular markers (**Table S1**) as above. After cell acquisition, the gates were drawn as shown in **Figure S4**.

### **Supplementary Tables and Figures**

***Supplementary Table 1.* Details of the reagents used in flow cytometry experiments*.***
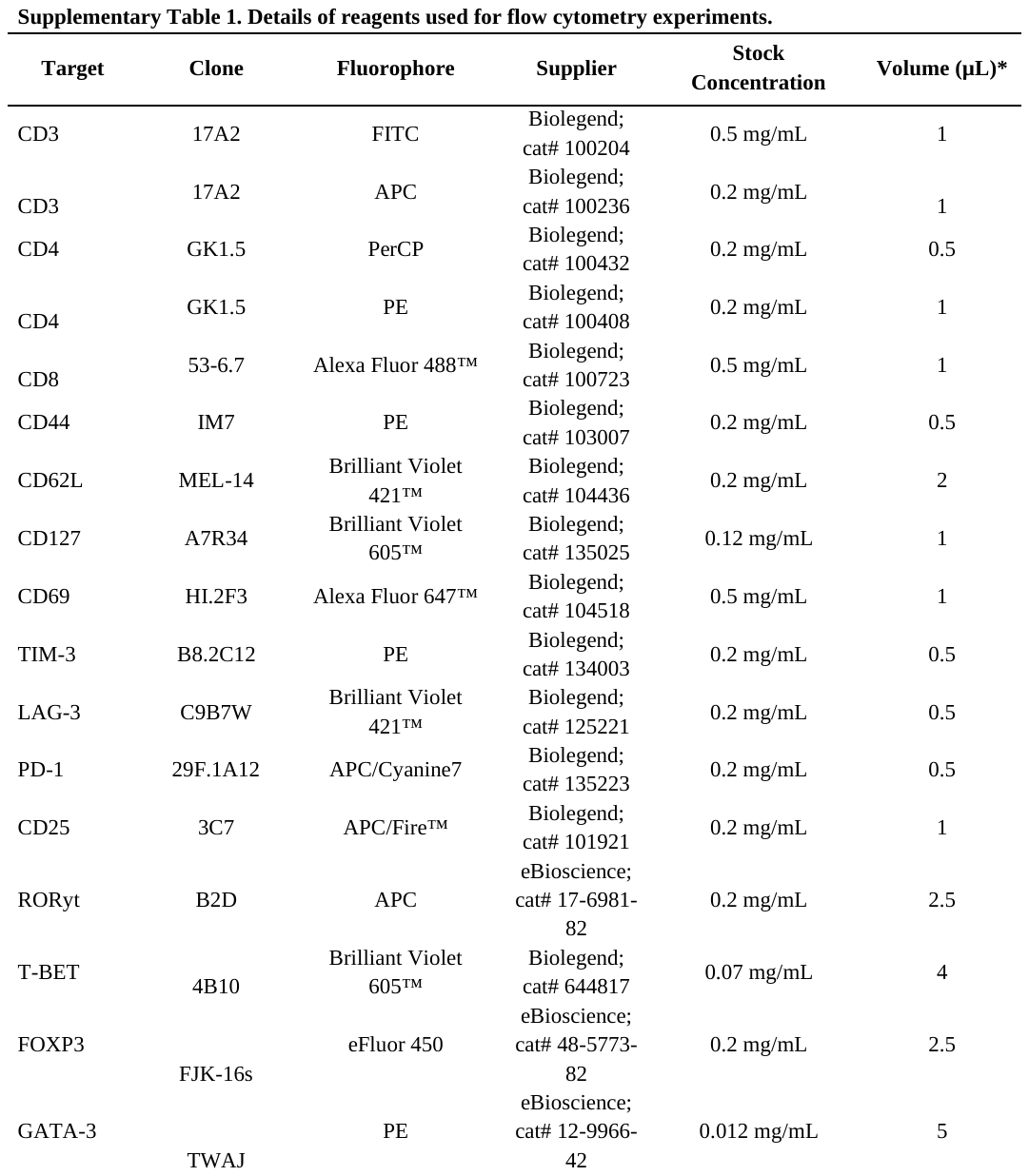

***Supplementary Table 2.* Markers used in the study and the phenotypes investigated using their combinations.**

| **Markers** | **Phenotype** |
| --- | --- |
| *CD4+ Phenotyping* | |
| CD3+CD4+ | CD4+ T Cells |
| CD3+CD4+GATA-3+ | Th2 Cells |
| CD3+CD4+T-bet+ | Th1 Cells |
| CD3+CD4+FoxP3+ | Treg Cells |
| CD3+CD4+RORγT+ | Th17 Cells |
| CD3+CD4+GATA-3+RORγT+ | Th2/Th17 Cells |
| CD3+CD4+T-bet+RORγT+ | Th1/Th17 Cells |
| CD3+CD4+FoxP3+RORγT+ | Treg/Th17 Cells |
| CD3+CD4+CD69+ | Early Activated T Cells |
| CD3+CD4+FoxP3-CD25+ | Late Activated T Cells |
| CD3+CD4+FoxP3-CD25+GATA-3+ | Th2 Late Activated Cells |
| CD3+CD4+FoxP3-CD25+T-bet+ | Th1 Late Activated Cells |
| CD3+CD4+FoxP3-CD25+RORγT+ | Th17 Late Activated Cells |
| CD3+CD4+PD-1+ | PD1+CD4+ T Cells |
| CD3+CD4+Tim-3+ | Tim-3+CD4+ T Cells |
| CD3+CD4+LAG-3+ | LAG-3+ CD4+ T Cells |
| CD3+CD4+PD-1+Tim-3+LAG-3+ | Terminally Exhausted CD4+ T Cells |
| CD3+CD4+CD62L+CD44+ | Central Memory CD4+ T Cells |
| CD3+CD4+CD62L-CD44+CD127+ | Effector Memory CD4+ T Cells |
| CD3+CD4+CD62L-CD44+CD127- | Effector CD4+ T Cells |
| *CD8+ Phenotyping* | |
| CD3+CD8+ | CD8+ T Cells |
| CD3+CD8+CD69+ | Early Activated CD8+ T Cells |
| CD3+CD8+CD25+ | Late Activated CD8+ T Cells |
| CD3+CD8+PD-1+ | PD1+CD8+ T Cells |
| CD3+CD8+Tim-3+ | Tim-3+CD8+ T Cells |
| CD3+CD8+LAG-3+ | LAG-3+CD8+ T Cells |
| CD3+CD8+PD-1+Tim-3+LAG-3+ | Terminally Exhausted CD8+ T Cells |
| CD3+CD8+CD62L+CD44+ | Central Memory CD8+ T Cells |
| CD3+CD8+CD62L-CD44+CD127+ | Effector Memory CD8+ T Cells |
| CD3+CD8+CD62L-CD44+CD127- | Effector CD8+ T Cells |
| CD3+CD8+KLRG1+ | Killer cell lectin-like receptor G1+ (KLRG1) CD8+ T Cells |
| CD3+CD8+KLRG1+CD127+ | Double-Positive Effector CD8+ T Cells |
| CD3+CD8+KLRG1+CD127- | Short-Lived Effector CD8+ T Cells |
| CD3+CD8+KLRG1-CD127+ | Memory Precursor Effector CD8+ T Cells |

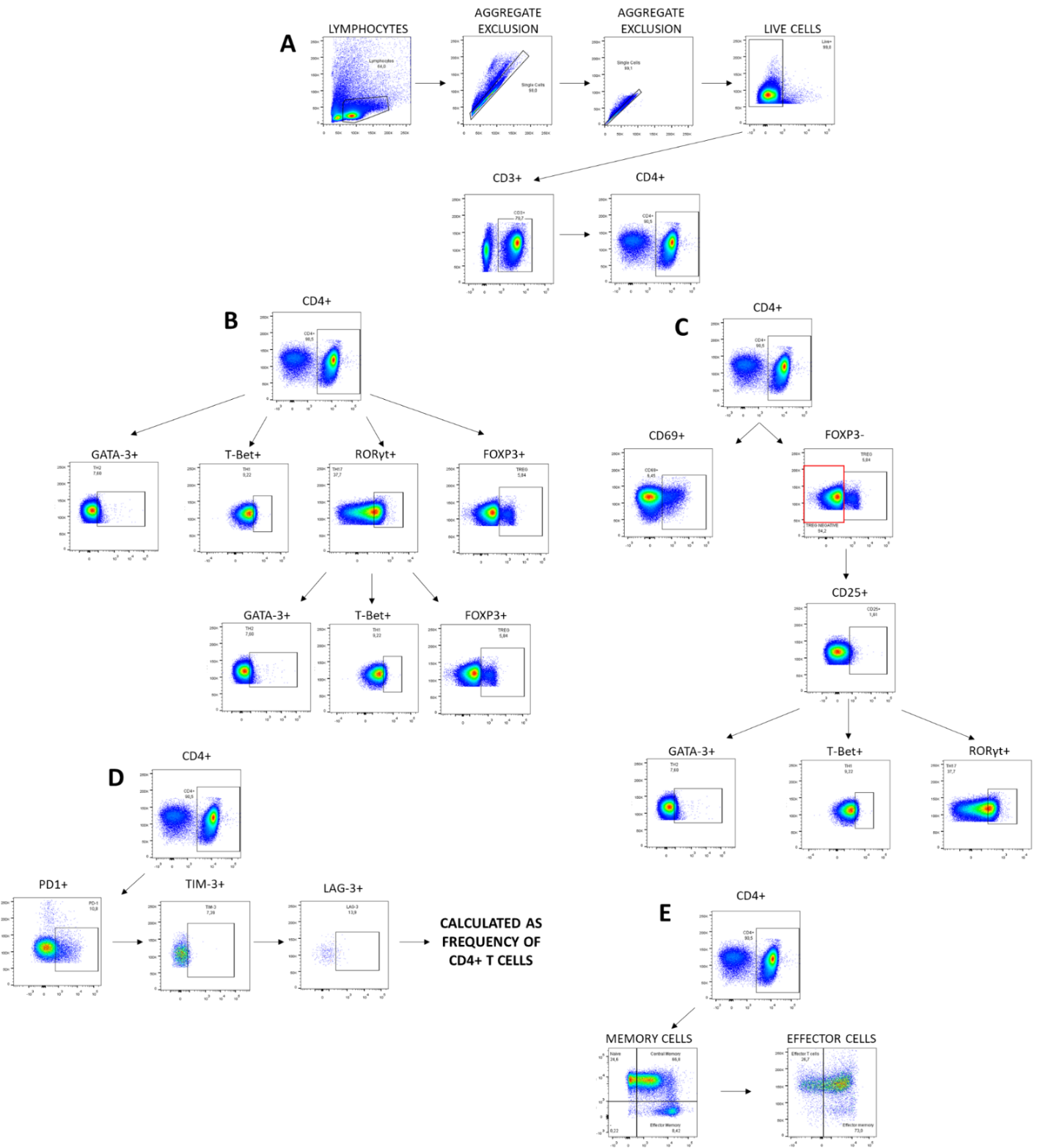

***Supplementary Figure 1. Gating strategies for CD4+ T-cell phenotyping experiments.* DR4tg mice were s**.c. injected with CFA and HomoCitJED or PBS **on day 0. Ten days later, splenocytes and dLNs were stained with various phenotypic markers, as indicated in A-E. A general gating strategy was applied to all panels to identify CD3+CD4+ T cells as in A, followed by additional gating to distinguish Th1/Th2/Th17/Treg subtypes in B, activated CD4+ T cells and their subtypes in C, exhausted CD4+ T cells in D, and effector/memory CD4+ T cells in E. Positive gates were determined based on FMO controls.**

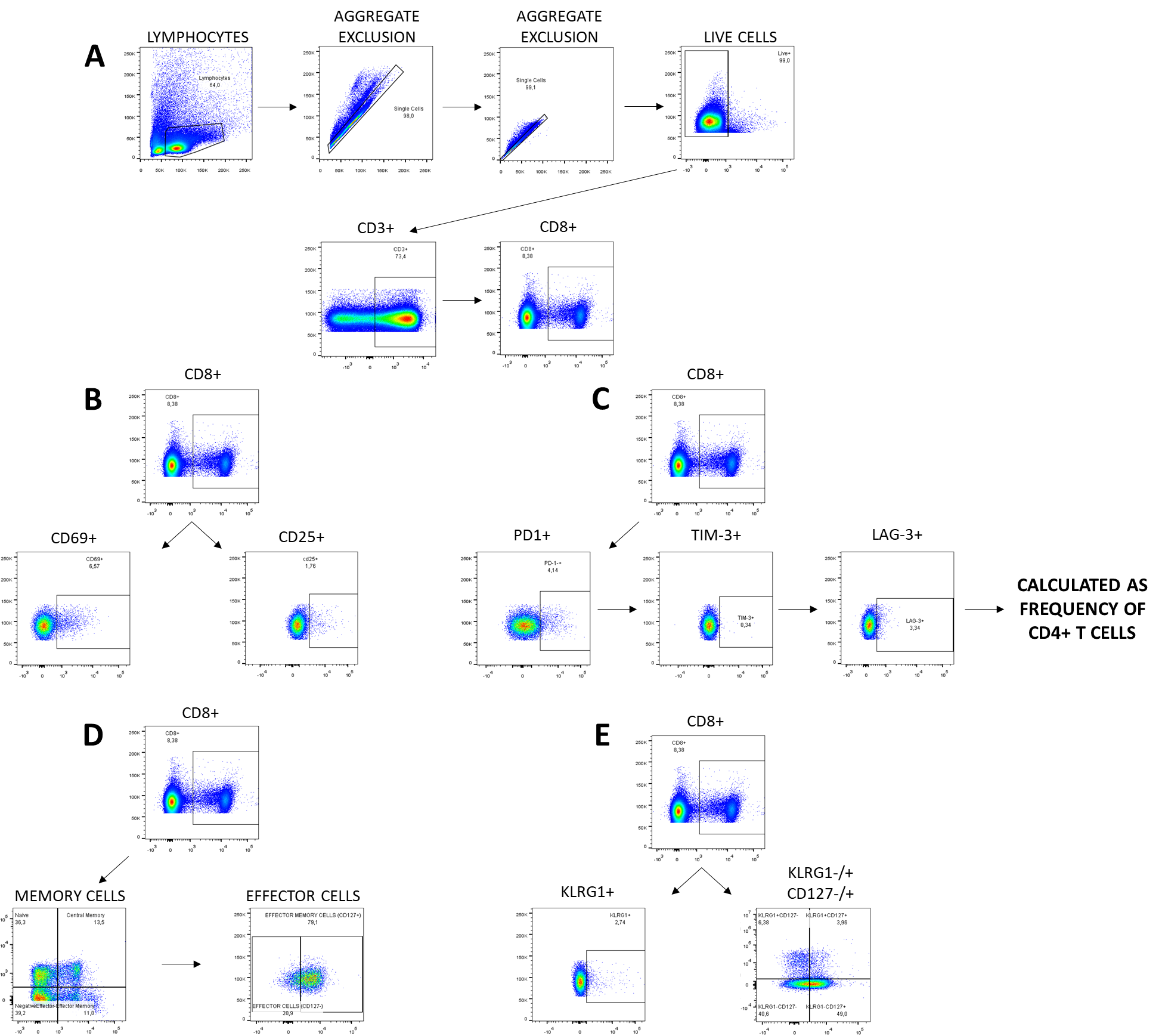

***Supplementary Figure 2. Gating strategies for CD8+ T-cell phenotyping experiments.*** DR4tg mice were s.c. injected with CFA and HomoCitJED or PBS on day 0. Ten days later, splenocytes and dLNs were stained with various phenotypic markers, as indicated in A-E. A general gating strategy was applied to identify CD3+CD8+ T cells as in **A**, followed by additional gating to distinguish activated CD8+ T cells as in **B**, exhausted T cells in **C**, effector/memory T cells in **D**, and KLRG1+/short/memory precursor effector T cells in **E**. Positive gates were determined using Fluorescence Minus One (FMO) controls.

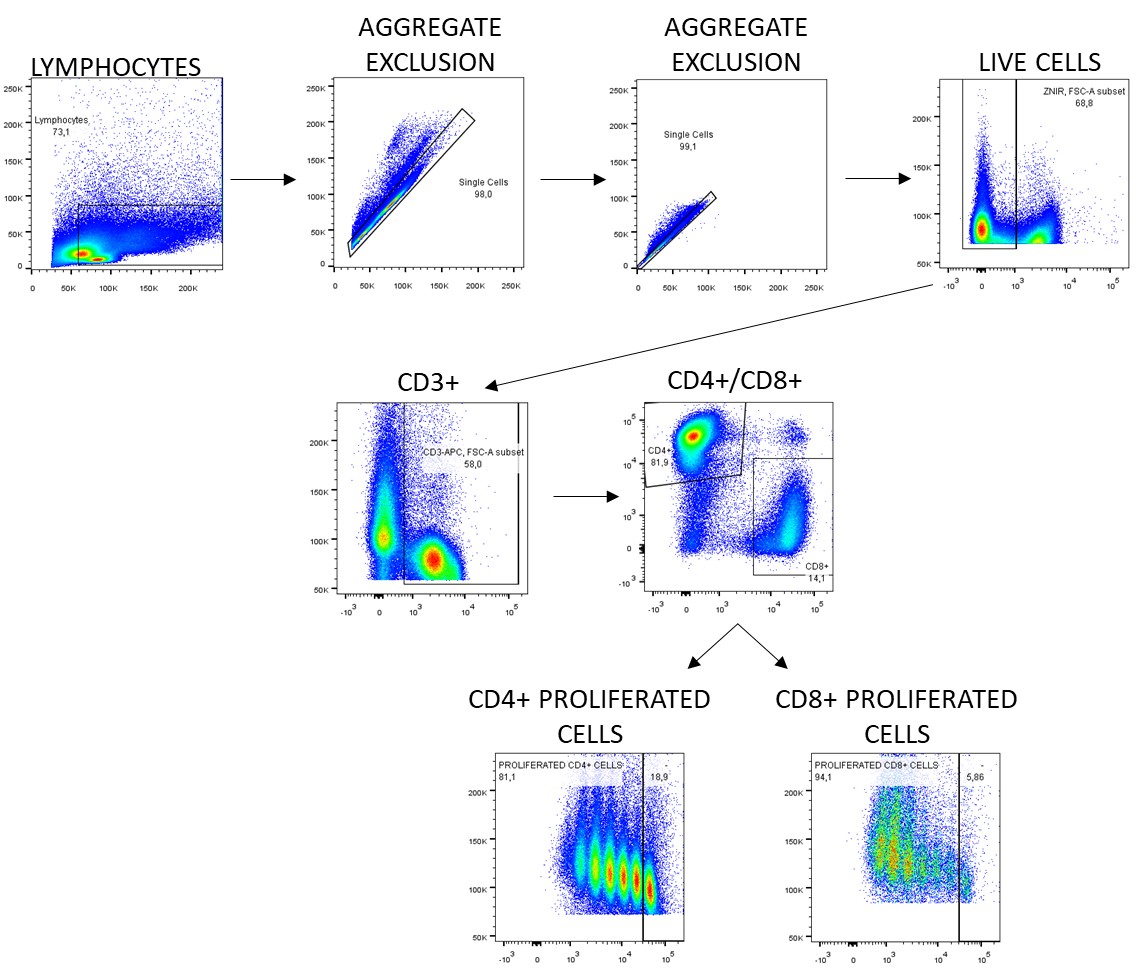

***Supplementary Figure 3. Gating strategies for proliferation experiments.*** Mice were s.c. injected with CFA and HomoCitJED or PBS on day 0. Ten days later, splenocytes and dLNs were stained with a proliferation dye, where each cell division reduces the dye's intensity, and cultured in the presence of 100 μg/mL of HomoCitJED, medium alone, or mitogens PMA (10 ng/mL) + ionomycin (100 ng/mL) as a positive control. After 72 hours of incubation, cells were washed and stained with various markers to detect the proliferation of T cells. A general gating strategy was applied to identify CD3+CD8+ or CD3+CD4+ T cells, followed by additional gating to measure proliferation. Proliferation gates were drawn based on medium alone samples and kept consistent for each condition in every mouse. Positive gates for other markers were determined using Fluorescence Minus One (FMO) controls.

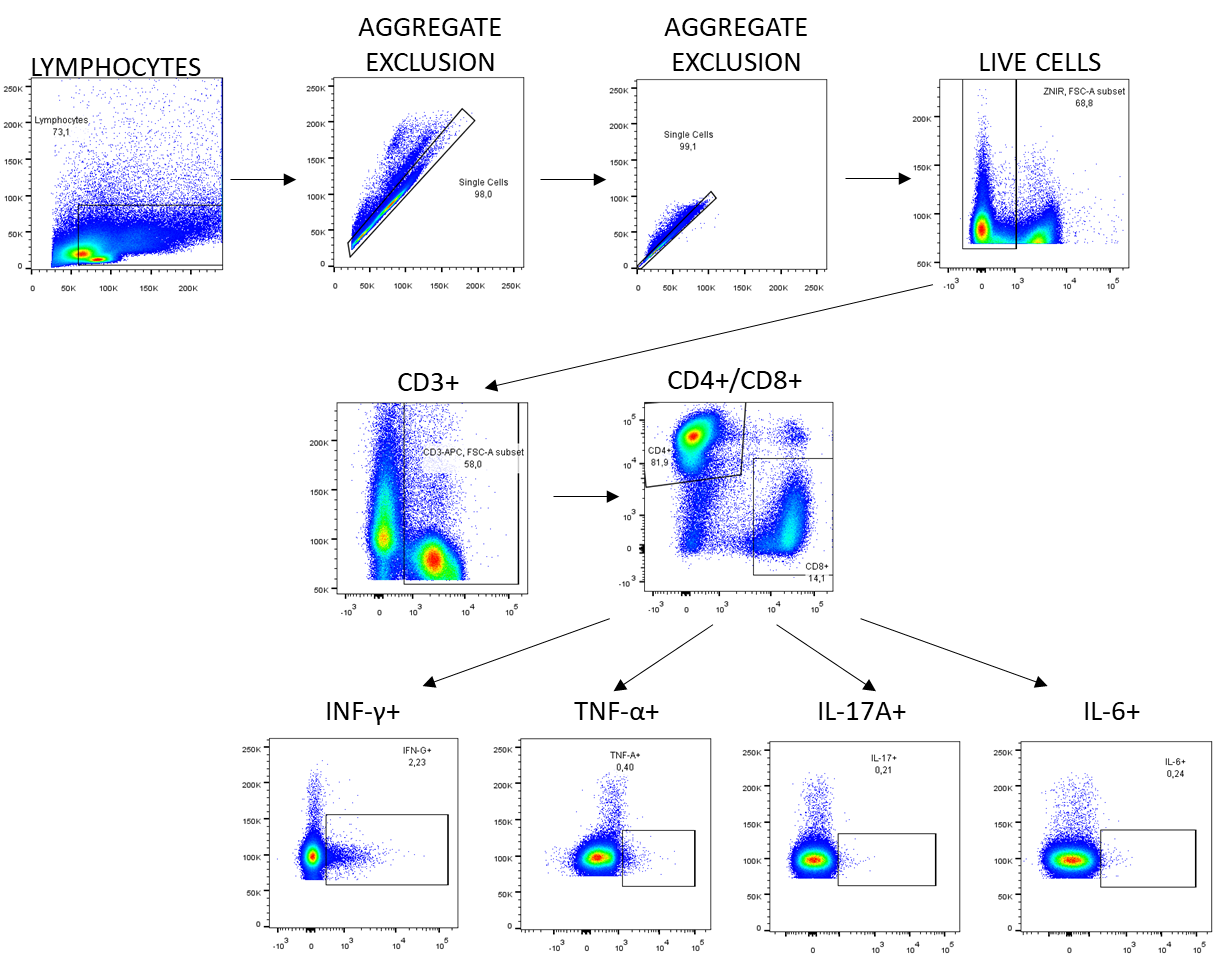

***Supplementary Figure 4. Gating strategy for intracellular cytokine staining experiments.*** Splenocytes and dLNs isolated from HomoCitJED+CFA- or PBS+CFA-s.c. injected mice on day 10 post-immunization were cultured in the presence of 100 μg/mL of HomoCitJED, medium alone, or the mitogens PMA (10 ng/mL) + ionomycin (100 ng/mL) as a positive control for 2 hours, after which the protein transport blocker Brefeldin A was added for the remaining 3 hours of incubation. After the overall 5-hour incubation, the production of TNF-α, IFN-γ, IL-6, and IL-17A by T cells was measured using flow cytometry. A general gating strategy was applied to identify CD3+CD8+ or CD3+CD4+ T cells, followed by additional gating to measure the cytokines of interest. Positive gates were determined using Fluorescence Minus One (FMO) controls and adjusted based on media alone controls.

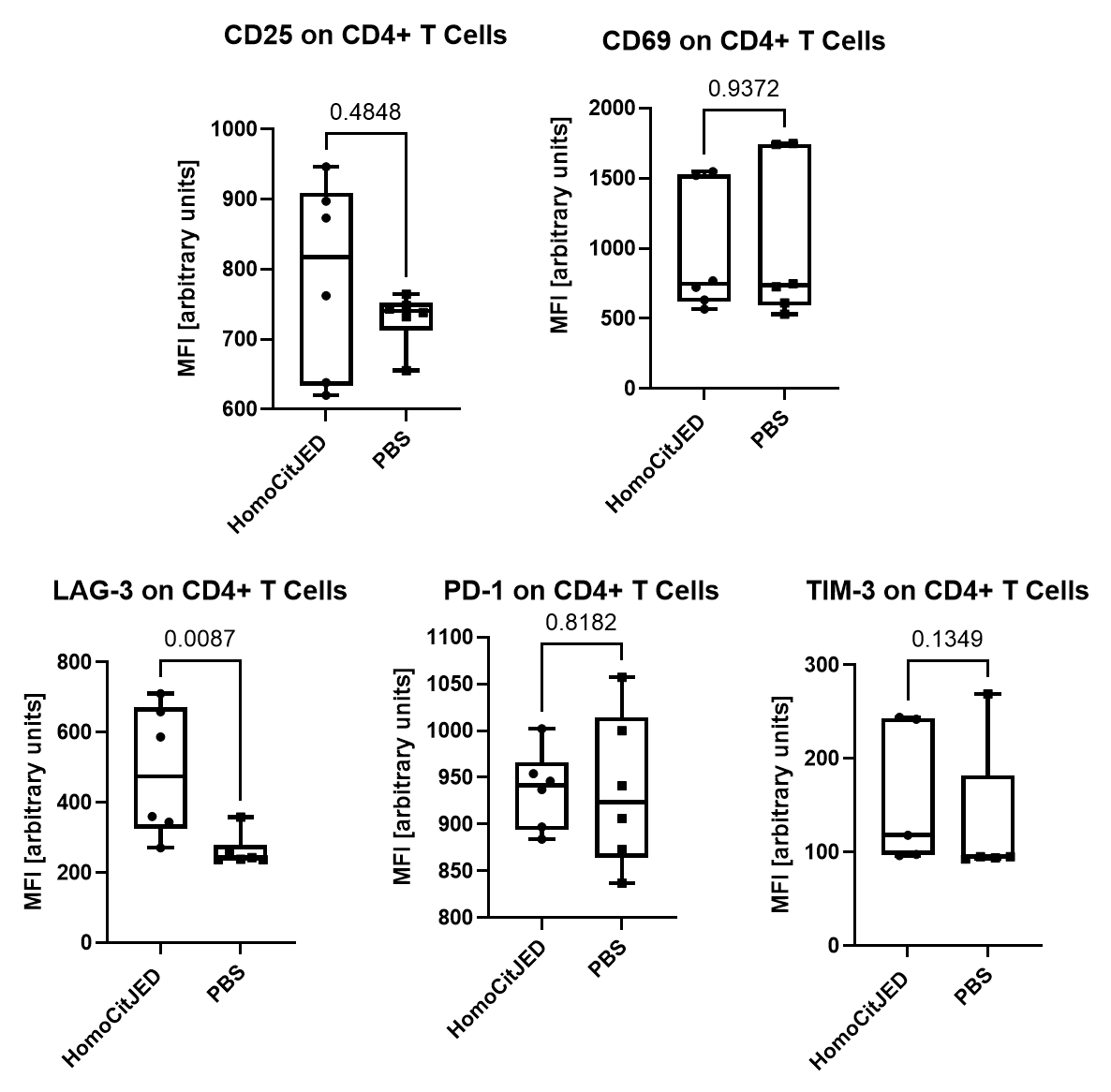

***Supplementary Figure 5. Activation and exhaustion marker expression on CD4+ T cells from DR4tg mice immunized with HomoCitJED.*** DR4tg mice were immunized s.c. with CFA and HomoCitJED or PBS on day 0; cells were isolated from dLNs on day 10 and analyzed by flow cytometry. Quantification of CD25, CD69, LAG-3, PD-1, and Tim-3 expression was performed using median fluorescence intensity (MFI) on CD4+ T cells. Graphs display the median and IQR for MFI; each symbol represents an individual mouse (n=6/group). Statistical comparisons were performed using the Mann–Whitney U test, with p < 0.05 considered significant.

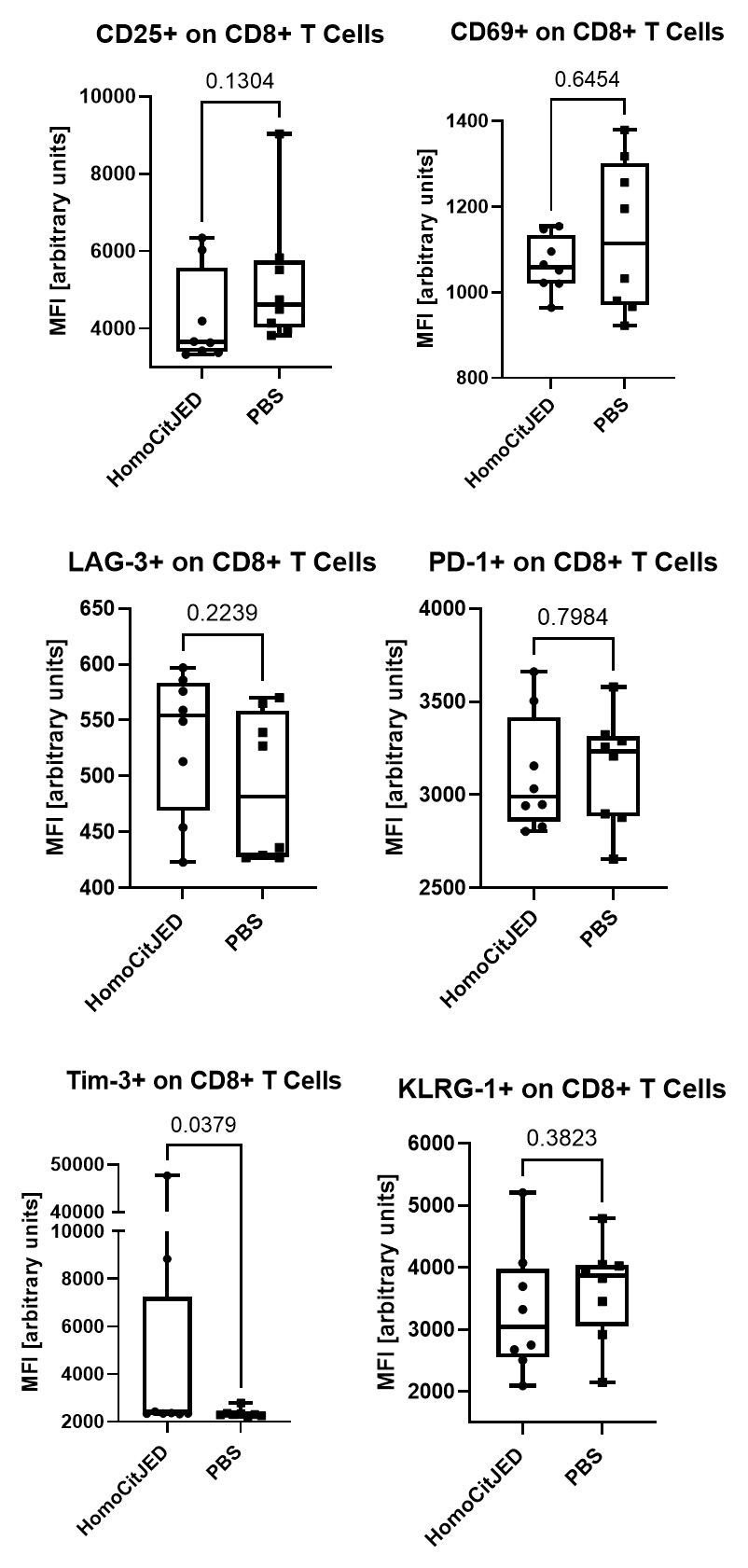

***Supplementary Figure 6. Activation and exhaustion marker expression on CD8+ T cells from DR4tg mice immunized with HomoCitJED.*** DR4tg mice were immunized s.c. with CFA and HomoCitJED or PBS on day 0; cells were isolated from dLNs on day 10 and analyzed by flow cytometry. Quantification of CD25, CD69, LAG-3, PD-1, Tim-3, and KLRG-1 expression was performed using median fluorescence intensity (MFI) on CD8+ T cells. Graphs display the median and IQR for MFI; each symbol represents an individual mouse (n=8/group). Statistical comparisons were performed using the Mann–Whitney U test, with p < 0.05 considered significant.

### **Supplementary Results**

### **2.1. Splenic CD4+ T‑cell responses to homocitrullinated peptides are enriched in FoxP3+RORγt+ and CD25+ T cells.**

To determine whether homocitrullinated peptides provoke systemic CD4+ T‑cell changes, we applied the same transcription factor and cell‑surface marker panels completed for dLNs to splenocytes harvested 10 days after s.c. injection with CFA and either HomoCitJED or PBS. Unlike the dLNs, splenic frequencies of canonical T‑bet+ Th1, RORγt+ Th17, GATA‑3+ Th2 and FoxP3+ Treg populations did not differ between HomoCitJED‑ and control mice (**Figure S7A**). Analysis of CD4+ T cells co‑expressing multiple lineage‑defining transcription factors showed that only FoxP3+RORγt+ (Treg/Th17) cells were expanded in HomoCitJED‑immunized animals compared with controls (1.5 % vs. 0.3 %, p=0.0152; **Figure S7B**).

Assessing functional activation markers revealed a modest but significant increase in CD25+ CD4+ T cells in the spleen of HomoCitJED‑immunized mice vs. controls (1.4 % vs 0.5 %, p = 0.0022; **Figure S7C**). No significant differences were observed between immunization groups for other activation, exhaustion or memory markers (**Figure S7C–E**).

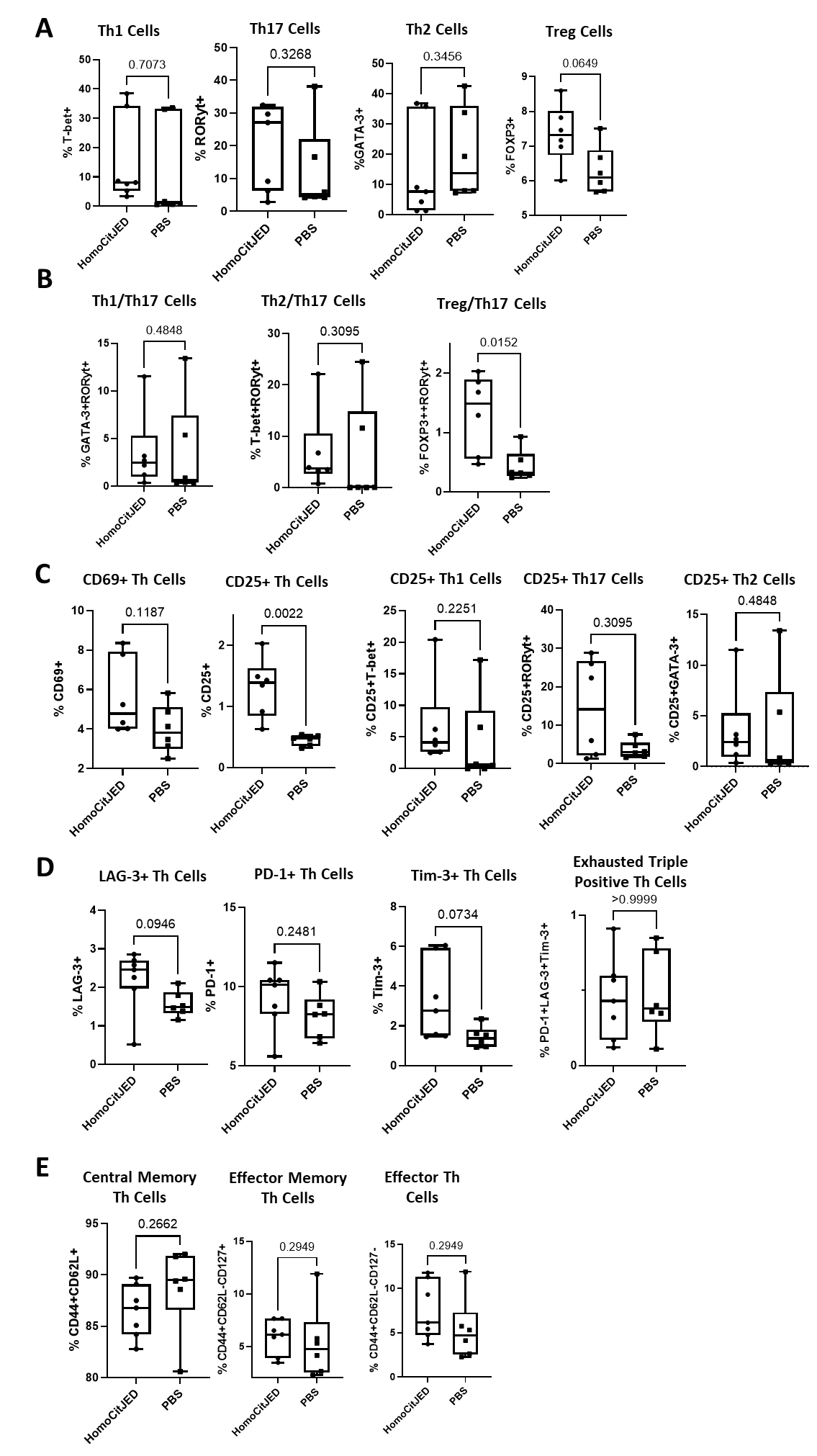

***Supplementary Figure 7. Th subtypes in splenocytes of DR4tg mice after immunization with HomoCitJED.*** DR4tg mice were s.c. injected with CFA and either HomoCitJED or PBS on day 0, and splenocytes were isolated on day 10 post-immunization. Flow cytometry was used to quantify T-bet+ Th1, RORγt+ Th17, GATA-3+ Th2, and FoxP3+ Treg cells, which are shown in **A**. Additional gating was performed to analyze double-positive subsets, including T-bet+RORγt+, GATA-3+RORγt+, and FoxP3+RORγt+ Th cells, as presented in **B**. Early activated CD69+ and late activated CD25+ CD4+ T cells were quantified, followed by further gating on CD25+ Th1, Th17, and Th2 subsets in **C**; FoxP3+ cells were excluded from the CD25+ activated cell analyses. The expression of exhaustion markers LAG-3, PD-1, and Tim-3, as well as triple-positive exhausted cells, is shown in **D**. Memory phenotypes, including CD44+CD62L+ central memory, CD44+CD62L-CD127+ effector memory, and CD44+CD62L-CD127- effector cells, were quantified and are presented in **E**. Values represent the percentage of total CD4+ T cells. Graphs display the median and IQR, with each symbol representing an individual mouse (n = 6–7/group). Statistical significance was determined using the Mann–Whitney U test, with p < 0.05 considered significant.

**2.2 Splenic CD8+ T‑cell phenotypes remain unchanged after HomoCitJED immunization**

The activation, exhaustion, and memory/effector‑differentiation profiles assessed in dLNs were applied in parallel to splenocytes collected 10 days after immunization. In contrast to the pronounced changes observed in dLNs, none of the analysed CD8+ T‑cell parameters differed between HomoCitJED‑ and control mice (**Figure S8**). Specifically:

1) **Early and late activation.** Frequencies of CD69+ and CD25+ CD8+ T cells were comparable between groups (**Figure S8A**).
2) **Checkpoint‑mediated exhaustion.** Expression of LAG‑3, PD‑1, and Tim‑3, individually or in triple‑positive terminally exhausted combinations, showed no significant differences (**Figure S8B**).
3) **KLRG1/CD127 and memory subsets.** Proportions of short‑lived effector cells (KLRG1+CD127-), double‑positive effector cells (KLRG1+CD127+), memory‑precursor effector cells (KLRG1-CD127+), and Tem, Tcm and Teff compartments were similar in HomoCitJED‑injected and control mice (**Figure S8C-D**).

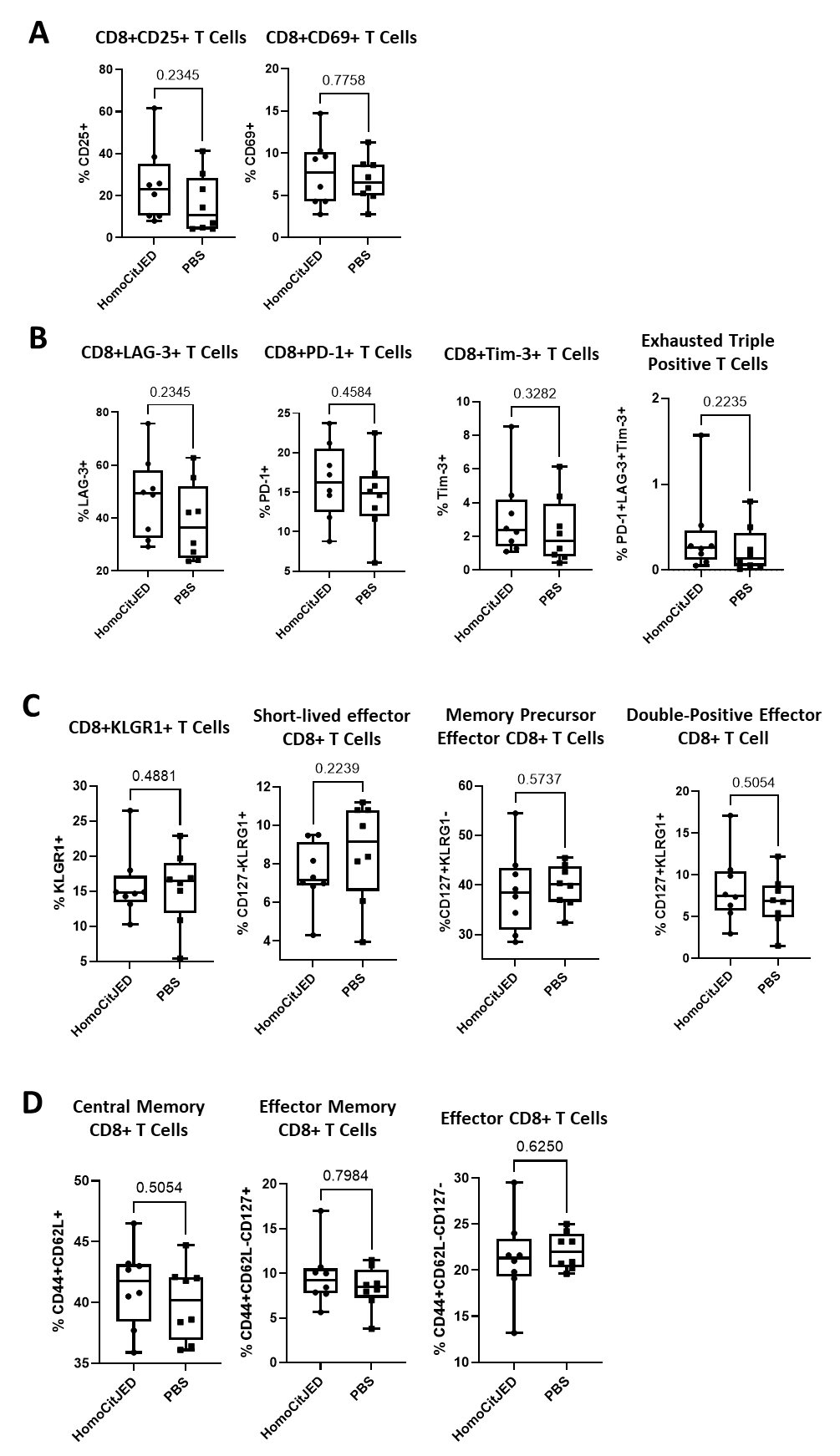

***Supplementary Figure 8. CD8+ T-cell phenotypes in splenocytes of DR4tg mice after immunization with HomoCitJED.*** DR4tg mice were s.c. injected with CFA and either HomoCitJED or PBS, and splenocytes were isolated on day 10 post-immunization. Flow cytometry was used to quantify early activated CD69+ and late activated CD25+ CD8+ T cells in **A**. The expression of exhaustion markers LAG-3, PD-1, and Tim-3, as well as triple-positive exhausted cells, is shown in **B**. Short-lived effectors (CD127-KLRG1+), memory precursors (CD127+KLRG1-), and double-positive effectors (CD127+KLRG1+) were quantified and are presented in **C**. Central memory (CD44+CD62L+), effector memory (CD44+CD62L-CD127+), and effector cells (CD44+CD62L-CD127-) were analyzed and are shown in **D**. Values represent the percentage of total CD8+ T cells. Graphs display the median and IQR, with each symbol representing an individual mouse (n=8/group). Statistical significance was determined using the Mann–Whitney U test, with p < 0.05 considered significant.

**2.3 Splenic T cells proliferate specifically in response to HomoCitJED**

We asked whether splenic T cells show a similar recall profile to the dLNs. HomoCitJED‑specific CD4+ proliferation was detected in 10/13 mice (77 %) versus 1/11 (9 %) controls, with median SI of 2.4 versus 1.1, respectively (p = 0.0003; **Figure S9**). Splenic CD8+ responses were observed in all HomoCitJED‑immunized mice (8/8, 100 %) but only 1/8 control animals (12.5 %), with SI of 2.9 versus 0.9, respectively (p = 0.0023; **Figure S9**).

The specificity control was splenocytes from HomoCitJED-immunized mice incubated with the lysine‑substituted peptide LysJED and revealed no detectable proliferation of either CD4+ or CD8+ splenic T cells (**Figure S10**). This confirms that the response is directed to homocitrulline rather than the peptide backbone.

Lastly, all mice responded to PMA + ionomycin mitogens, used as a positive control (data available upon request).

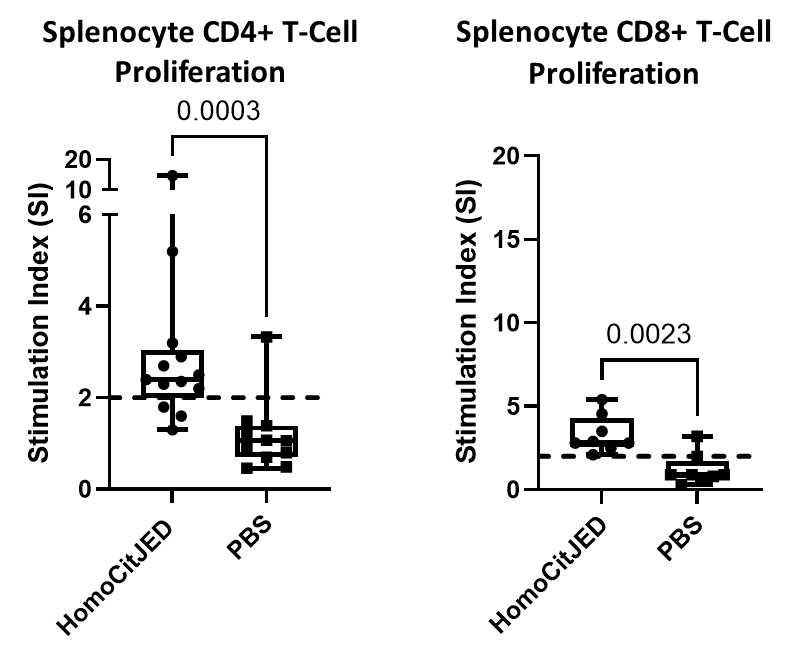

***Supplementary Figure 9. Proliferative responses of splenic T cells in HomoCitJED-immunized mice.***

Splenocytes from HomoCitJED+CFA- or PBS+CFA- s.c. injected mice were left untreated or cultured with 100 μg/mL HomoCitJED and analyzed by flow cytometry 72 hours later. Corresponding stimulation indices (SI) for CD4+ and CD8+ T-cell proliferations were determined by flow cytometry and are shown on the graphs. SI was calculated as the percentage of proliferating cells in peptide-stimulated samples divided by that in medium-only controls. A SI > 2.0 (indicated by a dashed line) was considered a positive proliferative response. Each symbol represents one mouse (n=8-13). Graphs show the median and IQR. Statistical significance was determined by the Mann–Whitney U test; p < 0.05 was considered significant.

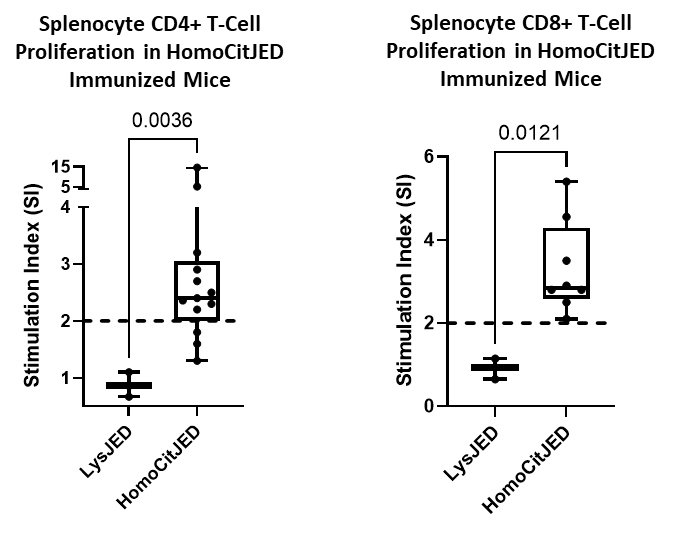

***Supplementary Figure 10. CD4+ and CD8+ T-cell proliferation to control peptide LysJED in mice immunized with HomoCitJED*.** Splenocytes isolated from HomoCitJED+CFA- s.c. injected mice were left untreated or cultured with 100 μg/mL HomoCitJED or control peptide LysJED and analyzed by flow cytometry 72 hours later. The proliferation of CD4+ and CD8+ T cells in the spleens, determined by flow cytometry, is shown as the median SI with IQR. SI was calculated as the percentage of proliferating cells in peptide-stimulated samples divided by that in medium-only controls. Stimulation indices greater than 2.0 (dashed line) were considered a positive proliferative response. Each symbol represents one mouse (n = 3-13/group). p < 0.05 by the Mann-Whitney U test was considered significant.

### **2.4. Homocitrullinated peptides induce pro‑inflammatory cytokine secretion in the splenocytes of DR4tg mice**

Given the central role of cytokines in inflammation observed in RA patients, we investigated the cytokine levels in the supernatants following a 48-hour splenocyte culture with HomoCitJED or media alone [3]. At this timepoint, we observed significantly higher levels of IL-17A, IL-2, TNF-α, and IFN-γ in the supernatants of mice immunized with homocitrullinated peptides compared with controls (**Figure S11**). In HomoCitJED-immunized mice versus controls, the highest concentration was observed for IL-17A (155 pg/mL vs. 0.4 pg/mL, p=0.0002), followed by TNF-α (25.4 pg/mL vs. 3.9 pg/mL, p=0.0070), IFN-γ (20.8 pg/mL vs. 0.005 pg/mL, p=0.0008), and IL-2 (10.1 pg/mL vs. 0.4 pg/mL, p=0.0009); no differences were detected in the concentration of IL-10 and IL-6 (**Figure S11**).

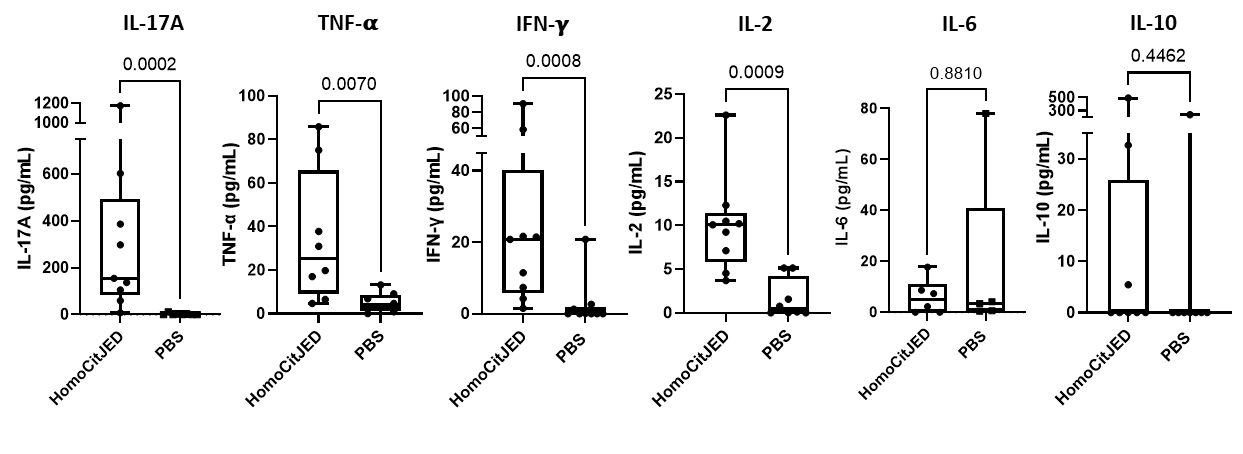

***Supplementary Figure 11. Cytokine concentrations in the supernatants of splenocytes treated ex vivo with HomoCitJED***. Splenocytes isolated from HomoCitJED+CFA- or PBS+CFA-injected mice on day 10 post-s.c. immunization were cultured in the presence of 100 μg/mL of HomoCitJED or medium alone. After a 48-hour incubation, the concentrations of IL-17A, TNF-α, IFN-γ, IL-2, and IL-6 were measured using ProQuantum immunoassays, and IL-10 was measured using ELISA. Cytokine concentrations were normalized by subtracting the concentration of the cytokine produced in the medium alone condition. Data are presented as the median and IQR (each symbol represents an individual mouse (n = 8–9/group)). A p-value < 0.05 by the Mann-Whitney U test was considered significant.

Because supernatant measurements do not identify the cellular source, we performed intra-cellular cytokine staining after a 5‑h *ex vivo* stimulation with HomoCitJED. Splenic CD4+ T cells from HomoCitJED‑immunized mice showed higher proportion of single‑positive IL‑6+ (3.85  vs 1.21, p = 0.0353) and IL‑17A+ (2.02 vs. 1.12, p = 0.0351) cells than controls (**Figure S12**). Dual IFN‑γ+IL‑17A+ CD4+ T cells were also enriched (1.82 % vs. 0.37 %, p = 0.0103) **(Figure S12**). No other CD4+ cytokine combinations and none of the CD8+ T‑cell cytokine subsets differed between the groups (**Figures S12-S13**). PMA + ionomycin elicited robust production of measured cytokines in both cohorts, validating the assay (data available upon request).

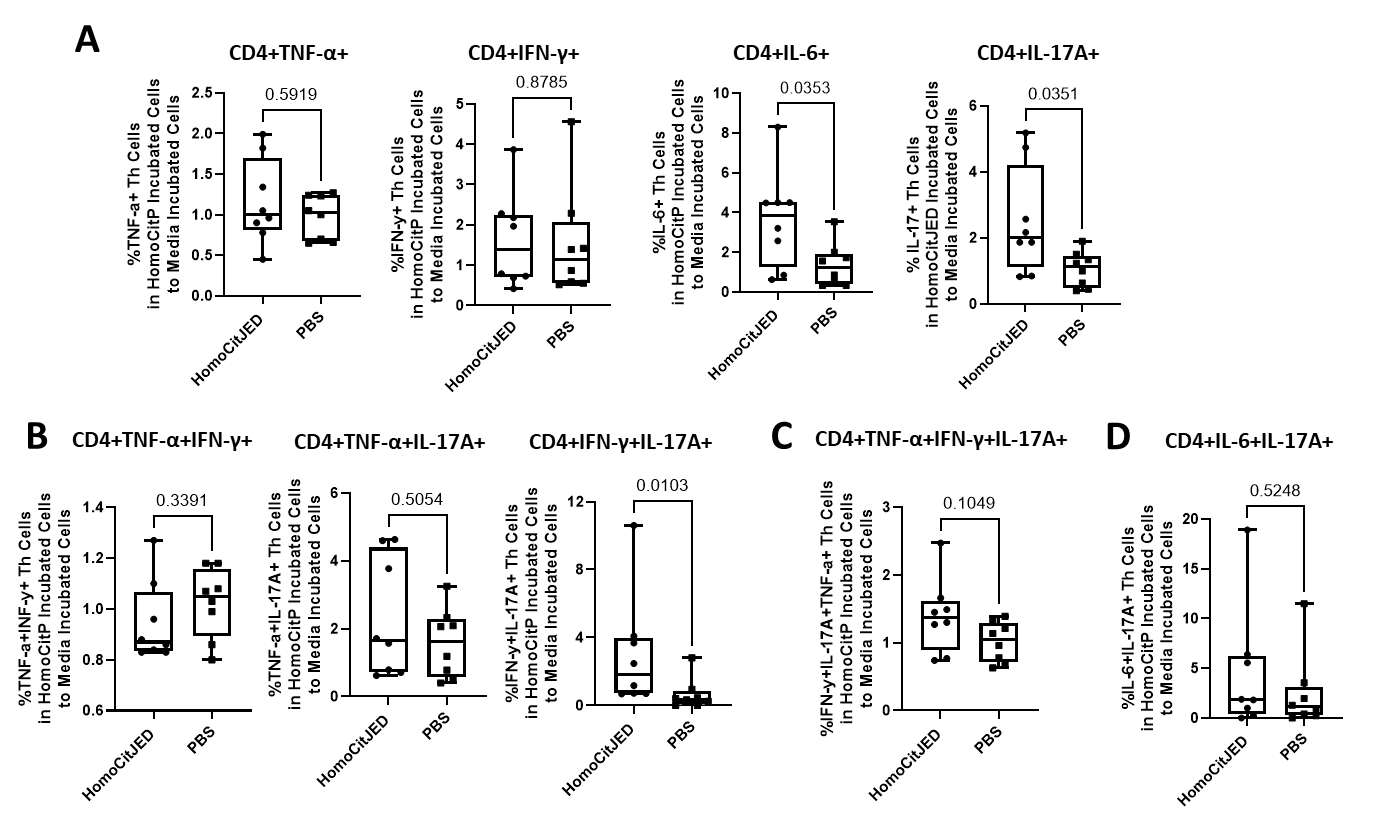

***Supplementary Figure 12. Cytokine profiles of CD4+ T cells from spleens after ex vivo stimulation with HomoCitJED***. DR4tg mice were s.c. injected with CFA and HomoCitJED or PBS and euthanized 10 days post-s.c. injection. Splenocytes were cultured with 100 µg/mL HomoCitJED or medium alone for 5 h. Intracellular staining for CD4+ T cells expressing TNF-α, IFN-γ, IL-6 and IL-17A was analyzed by flow cytometry. Cells producing a single cytokine are shown in **A**. Cells producing Th1-type cytokines or both a Th1 and a Th17 cytokine are represented in **B**. Cells producing three cytokines characteristic of Th1/Th17 responses are depicted in **C**. Cells co-expressing IL-17A with IL-6, consistent with Th17/Th2 plasticity are quantified in **D**. Cytokine production is expressed as the ratio of cells stimulated with HomoCitJED to those cultured in media alone. Graphs display the median and IQR; each symbol represents an individual mouse (n = 8/group). Statistical significance was determined by the Mann-Whitney U test; p < 0.05 was considered significant.

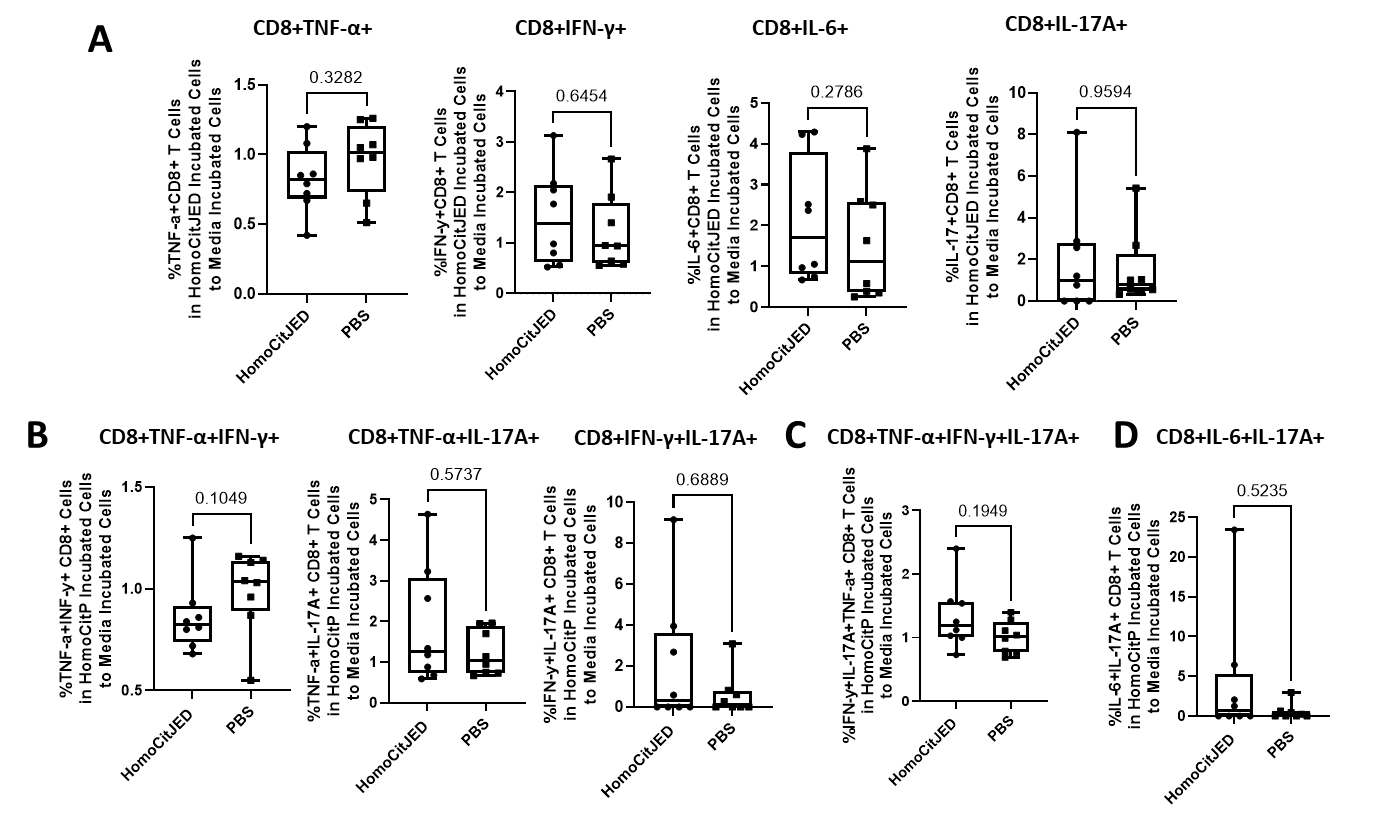

***Supplementary Figure 13. Cytokine profiles of CD8+T cells (Tc subsets) from dLNs after ex vivo stimulation with HomoCitJED***. DR4tg mice were s.c. injected with CFA and HomoCitJED or PBS and euthanized on day 10 post-injection. Splenocytes were cultured with 100 µg/mL HomoCitJED or medium alone for 5 h. Intracellular staining for CD8+ T cells expressing TNF-α, IFN-γ, IL-6 and IL-17A was analyzed by flow cytometry. Cells producing a single cytokine are shown in **A**. Cells producing Tc1-type cytokines or both a Tc1 and a Tc17 cytokine are represented in **B**. Cells producing three cytokines associated with Tc1/Tc17 responses are depicted in **C**. Cells co-expressing IL-17A with IL-6, consistent with Tc17/Tc2 plasticity are quantified in **D**. Cytokine production is expressed as the ratio of cells stimulated with HomoCitJED to those cultured in medium alone. Graphs display the median and IQR; each symbol represents an individual mouse (n = 8/group). Statistical significance was determined by the Mann-Whitney U test; p < 0.05 was considered significant.
